## Supplementary material for "Using informative priors to account for identifiability issues in occupancy models with identification errors": latex files: _FINAL.tex

\documentclass[PCJ,Unicode,screen]{cedram}

\usepackage[natbib=true, mincitenames=1, maxcitenames=2, maxbibnames=20,minbibnames=20, uniquelist=false, uniquename=false, style=pcj, url=false, defernums=true]{biblatex} % pour la biblio % backend=biber, 
\usepackage{setspace}
% This is to correct wrong references sorting, see sample .bib file.
\providecommand*{\noopsort}[1]{}

\newcommand{\defeq}{\vcentcolon=}

\usepackage{mathtools}
\usepackage{chngcntr}
% for PCI badge
\usepackage{tikz}
% to put it on the title page
\usepackage{atbegshi} 
%\usepackage{titling}
\graphicspath{ {./graphs/} }

\newcommand{\addPCIbadge}{
    \ifnum\value{page}=1
    \begin{tikzpicture}[overlay] %remember picture
       \hypersetup{hidelinks}
       \node[anchor=north west, yshift=3.5cm, xshift=14.5cm][rotate=-20]
       % \node[anchor=north west,yshift=60pt,xshift=-20pt]
           {\href{https://doi.org/10.24072/pci.ecology.100699}
           {\includegraphics[height=45mm]{badge_PCI_Ecology.png}}} ;
    \end{tikzpicture}
    \fi
}


%%%%%%%%%%%%%%%%%%%%%%%%%%%%%%%%%%%%%%%%%%%%%%%%%%%
 % Declare your BibTeX file here (keep the .bib extension):
\addbibresource{references2.bib} %_maBiblio.bib

\title{Using informative priors to account for identifiability issues in occupancy models with identification errors}

\DOI{10.1101/2024.05.07.592917}
%\PCIcorresp{}

\author{\firstname{Célian} \lastname{Monchy}}
\address{CEFE, Univ Montpellier, CNRS, EPHE, IRD, Montpellier, France}

\author{\firstname{Marie-Pierre} \lastname{Etienne}\IsEqualContrib}
\address{IRMAR - Institut de Recherche Mathématique de Rennes}
\author{\firstname{Olivier} \lastname{Gimenez}\IsEqualContrib}
\address{CEFE, Univ Montpellier, CNRS, EPHE, IRD, Montpellier, France}
 
\keywords{Bayesian modelling, camera traps, environmental DNA, false-positive, identifiability,  informative priors, misidentification, non-invasive sampling, species occupancy}

\begin{abstract}
 Non-invasive monitoring techniques like camera traps, autonomous recording units and environmental DNA are increasingly used to collect data for understanding species distribution. These methods have prompted the development of statistical models to suit specific sampling designs and get reliable ecological inferences.

Site occupancy models estimate species occurrence patterns, accounting for the possibility  that the target species may be present but unobserved. Here, two key processes are crucial: detection, when a species leaves signs of its presence, and identification where these signs are accurately recognized. While both processes are prone to error in general, wrong identifications are often considered as negligible with in situ observations. When applied to passive bio-monitoring data, characterized by datasets requiring automated processing, this second source of error can no longer be ignored as misclassifications at both steps can lead to significant biases in ecological estimates. Several model extensions have been proposed to address these potential errors.

We propose an extended occupancy model that accounts for the identification process in addition to detection. Similar to other recent attempts to account for false positives, our model may suffer from  identifiability issues, which usually require another source of data with perfect identification to resolve them. As an alternative when such data are unavailable, we propose leveraging existing knowledge of  the identification process within a Bayesian framework by incorporating this knowledge through an informative prior. Through simulations, we compare different prior choices that encode varying levels of information, ranging from cases where no prior knowledge is available, to instances with accurate metrics on the performance of the identification, and scenarios based on generally accepted assumptions. We demonstrate that, compared to using a default prior, integrating information about the identification process as a prior reduces bias in parameter estimates. Overall, our approach mitigates identifiability issues, reduces estimation bias, and minimizes data requirements.

In conclusion, we provide a statistical method applicable to various monitoring designs, such as camera trap, bioacoustics, or eDNA surveys, alongside non-invasive sampling technologies, to produce ecological outcomes that inform conservation decisions.
\end{abstract}

\begin{document}

\thispagestyle{empty}
\nolinenumbers
\AtBeginShipout{\AtBeginShipoutAddToBox{\addPCIbadge}}

\maketitle

\onehalfspacing

%\clearpage
%\setcounter{page}{1}

\section*{Introduction}

A primary objective for ecologists and conservation scientists is to understand how populations and communities are distributed across space and time. 
Monitoring animal species, plants, and even pathogens typically involves collecting data on their presence, and ideally, their absence, in order to evaluate their distribution area. Occupancy models have been developed by MacKenzie et al. (2002, see also \cite{tyre_improving_2003}) to account for potential undetected  presence. These models estimate the proportion of sites occupied by a species while accounting for the imperfect detection of the species during field surveys \parencite{mackenzie_estimating_2002}. Since a single visit is not sufficient to distinguish between a  present but undetected species and its true absence from a site, \textcite{mackenzie_estimating_2002} showed that repeated visits to the same site enable the estimation of the false-negative error rate, defined as the probability that a species present at a site remains undetected during a visit.
Over the last decade, the development of new, non-invasive monitoring techniques such as camera traps (e.g. \cite{parsons_occupancy_2017, hofmeester_framing_2019}), autonomous acoustic recording units (e.g. \cite{wrege_acoustic_2017, shonfield_autonomous_2017}) and environmental DNA sampling (e.g. \cite{griffin_modelling_2020,da_silva_neto_integrating_2020}) has deeply changed data collection for biodiversity monitoring. The integration of passive sensor technologies into conservation projects is expanding, driven by technical improvements that facilitate the efficient monitoring of multiple species, including cryptic taxa, across large areas and challenging environments \parencite{burton_review_2015}.
However, these emerging methods are not exempt from imperfect detection. Indeed, certain discrete taxa may remain silent, do not trigger camera traps, or leave minimal detectable traces \parencite{belmont_new_2022, goldman_integrating_2023}, so it remains essential to consider the probability of detecting them, regardless of the observation method used.

Within the context of sensor-based assessment method, data are massive and need to be processed before being analyzed. In particular, this involves identifying the taxon of interest in a large amount of collected data, either manually by operators \parencite{swanson_snapshot_2015, welbourne_effectiveness_2015}, through automated deep learning algorithms \parencite{tabak_machine_2019, duggan_approach_2021}, or a combination of both  \parencite{augustine_towards_2023, campos-cerqueira_improving_2016}. This step raises many statistical challenges \parencite{hartig_novel_2024}. 
For images and acoustic data, combining manual and automated processing helps to control classification errors; such as  misidentifying one species as another \parencite{barre_accounting_2019}. 
Similarly, environmental DNA studies also generate large datasets from which presence data must be extracted \parencite{hunter_environmental_2015, thomsen_monitoring_2012, schmidt_site_2013}. Detecting an organism's presence from its DNA in the environment is subject to various sources of variability, including the molecular techniques employed, laboratory procedures, and the amount of DNA collected  \parencite{willoughby_importance_2016, doi_evaluation_2019}.  Despite the sensitivity of molecular techniques, once data are processed, distinguishing between real absences and those resulting from poor sampling or identification errors remains challenging \parencite{goldberg_critical_2016}. Thus, it is essential to consider both mis-identification and mis-detection in eDNA surveys.  
In eco-epidemiology studies, site occupancy models are used to estimate the occurrence of pathogens responsible for wildlife diseases within a sample unit, providing insights into spatial patterns and disease dynamics \parencite{mcclintock_seeking_2010}. The challenge for wildlife disease surveys is similar to that in camera-trapping for conservation, as both involve estimating occupancy parameters based on imperfect diagnostic tests \parencite{thompson_use_2007, mcclintock_seeking_2010, lachish_site-occupancy_2012}.

The challenges of studies based on new biomonitoring technologies stem from the sequential nature of the detection and identification processes, each of which introduces two types of errors. A false-negative mis-identification occurs when a species is detected (e.g., the camera is triggered) but not correctly identified. Conversely, a false-positive mis-identification occurs when a species is not detected, but an error in data processing leads to its accidental identification. This two-step process increases the likelihood of errors in eDNA or sensor-based studies, compared to conventional surveys \parencite{hartig_novel_2024}.
Failure to account for these identification errors can result in biased estimates of the actual proportion of occupied sites \parencite{mackenzie_estimating_2002, tyre_improving_2003, spiers_estimating_2022}. 
The standard site occupancy model accounts for false-negative errors by estimating the probability of imperfect detection, but it does not account for the possibility of false-positive detections, where a species is incorrectly identified at a site it does not occupy. False-positive errors, if unaddressed, can lead to overestimating occupancy probability \parencite{royle_generalized_2006, mcclintock_unmodeled_2010, miller_improving_2011}. Consequently, several authors have proposed extending MacKenzie's site occupancy model by accounting for false detection, although these extensions face identifiability issues \parencite{chambert_modeling_2015} often resolved by incorporating additional data sources, including one without errors. 
For example, \textcite{miller_improving_2011} proposed a multiple detection state model in which both certain and ambiguous data are used at each site.  Building on this,  \textcite{chambert_modeling_2015} introduced the concept of “reference sites” exempt from detection error, and \textcite{mckibben_model_2023} revisited the notion of detection ambiguity introduced by \textcite{miller_improving_2011} by scoring observer confidence levels.

While these studies offer solutions for addressing detection errors, especially false positives, they rely on the integration of different data sources, which represents a strong constraint that cannot always be met. Indeed, great logistics and human efforts are often needed to design sampling protocols, collect and/or verify data, and to finally get several sources of data with some of them guaranteed to be error free. Although error-free data are rarely available, some knowledge about the reliability of the identification process may still be accessible (e.g., expert beliefs, calibration experiments or performance metrics). In this case, eliciting informative prior distribution may be an alternative to the combination of several sources of data \parencite{guillera-arroita_dealing_2017, cruickshank_quantifying_2019}. The use of Bayesian statistics allows the integration of information through informative prior, which has been shown to increase confidence in the results \parencite{mccarthy_profiting_2005, choy_elicitation_2009}. In occupancy studies with sparse data, a precise choice of priors influences trend occupancy estimates \parencite{outhwaite_prior_2018}. However, those informative priors must be chosen carefully, in accordance with the available knowledge, otherwise the parameter estimates could be biased \parencite{morris_neglected_2015}. Here, we propose a hierarchical model that builds on the classical occupancy model to account for identification errors across different types of data.
We first provide a probabilistic description of the model, discuss the limitations of a frequentist approach for fitting this model, and then propose to overcome these limitations using a Bayesian framework that allows incorporating available information through informative priors.  Through simulations, we compare the effectiveness of the different approaches. 

\section*{Model Description}
\subsection*{Standard Occupancy model}

Detection and non-detection data on a species are collected from $S$ sites, visited $J$ times. These repeated visits help differentiate between sites where the species is truly absent and those where the species is present but not detected. In the hierarchical formulation of the occupancy model \parencite{mackenzie_estimating_2002} the latent occupancy state of a site $i$ is a Bernoulli distributed random variable of parameter $\psi$, hence the species is present on a site $i$ ($Z_i=1$) with a probability $\psi$: 
\begin{align}
    \label{eq:standardmodel}
    Z_{i} &\overset{i.i.d}{\sim}Bernoulli(\psi) \\ \nonumber
    Y_{ij} | Z_i &\overset{i.i.d}{\sim} Bernoulli(Z_{i} \times p)
\end{align} 
Furthermore, it is assumed that species presence at one site is independent of its presence at other sites, meaning that $Z_i$ (with $i$ from 1 to $S$) are independent.
Given the species is present at site $i$, $Y_{ij}$ represents the detection state during visit $j$. It follows a Bernoulli distribution with parameter $p$, such as the species may be detected with a probability $p$ during the $j^{th}$ visit on the occupied site $i$, and missed with probability $1-p$. 
In this model, each visit is considered as an observation, the species being detected or not. Conditionally on the presence ($Z_i=1$), the history of detection is a set of independent observations for a site, represented by a vector of detections (1) and non-detections (0). 

While this model is appropriate for traditional field observations, it can be adapted according to the monitoring method. For some species, passive biomonitoring techniques offer a cost-effective alternative to field observations, but introduce new challenges. Unlike direct field observations, sensor data must be processed to determine species presence, and this introduces potential errors in detection history, including false positives, which are not accounted for in the standard occupancy model.

\subsection*{Extended model to identification level}

To address these challenges, we extend the original model by introducing an additional identification process that accounts for potential errors in species identification. This step is particularly important when working with data where species identification can be ambiguous. 

In this extended model, the potential detection becomes a latent variable $Y_{ij}$ and we add a second layer to account for potential error  in the identification process: an observation may correspond to a record (acoustic or image) where the species is identified (either correctly or incorrectly). Detection, however remains an unknown variable, referring to the sensor triggering and capturing the species' presence. In some cases, where the quality of the recorded file is too poor or for species difficult to differentiate, the species may be detected but not correctly identified  \parencite{findlay_component_2020}. Thus it is impossible to deduce the detection state from the record alone.

To formalize this, we denote $W_{ij}$ as the species identification at site $i$ on visit $j$. $W_{ij}$ equals 1 if the species is identified and 0 otherwise. The identification process is imperfect and suffers from two types of error related to the detection or non-detection of the species, each with its own probability (Fig. \ref{fig:tree}). The probability to identify the species in the $j^{th}$ visit from site $i$ if it has been detected is $w_A$, and it is equivalent to the probability of correctly identify the detected species. This is related to the true positive probability, also known as sensitivity or \textit{recall}. Otherwise, the probability to falsely identify the species while it has not been detected is $1-w_B$, usually referred to as the false positive rate (also known as \textit{fall-out}), and corresponding to the probability of associating an observation to the wrong species. 

In contrast to the standard model from \textcite{mackenzie_estimating_2002}, where the identification errors are not considered, assuming that $w_A=1$ and $w_B=1$, this extended model explicitly accounts for the possibility of false identifications. In other words, the probability of failing to identify a species that has been detected is zero, as is the probability of confusing an undetected species with a detected one.

Given this extended framework, the conditional probability of identifying a species $W_{ij}=1$ given that it is detected or not is written as:

\begin{gather}
\label{eq:model_w}
    W_{ij} |Y_{ij} =  
        \left \{
          \begin{array}{ll}
            W_{ij} | (Y_{ij}=0) \sim Bernoulli(1 - w_{B,ij}) \\
            W_{ij} | (Y_{ij}=1) \sim Bernoulli(w_{A,ij})
          \end{array}
        \right.
\end{gather}

In this hierarchical model, $Z_i$ and $Y_{ij}$ are latent variables respectively related to occupancy state and detection state of the target species at site $i$ during visit $j$, and where $W_{ij}$ is the observation data related to identification (Fig. \ref{fig:tree}).

\begin{figure}[h!]
    \centering
    \begin{tikzpicture}[level 1/.style={sibling distance=90mm},
    level 2/.style={sibling distance=50mm}, level 3/.style={sibling distance=20mm}]
\node []{} %$\bullet$
  child { node [circle] {$Z_i=1$}
        child { node {$Y_{ij}=1$}
               child { node [draw] {$W_{ij}=1$} 
	        	  edge from parent [->] node [left] {$w_A$} 
        		}  
        		child { node [draw] {$W_{ij}=0$} 
	        	  edge from parent [->] node [right] {$1-w_A$} 
				}  
        	  edge from parent [->] node [left] {$p$} 
   			}
        child { node {$Y_{ij}=0$}
        		child { node [draw] {$W_{ij}=1$} 
	        	  edge from parent [->] node [left] {$1-w_B$} 
        		}  
        		child { node [draw] {$W_{ij}=0$} 
	        	  edge from parent [->] node [right] {$w_B$} 
				}  
        	  edge from parent [->] node [right] {$1-p$} 
   			}
     edge from parent [->] node [left] {$\psi\quad $} 
     }
  child { node [circle] {$Z_i = 0$}
        child { node {$Y_{ij}=0$}
        		child { node [draw]{$W_{ij}=1$} 
	        	  edge from parent [->] node [left] {$1-w_B$} 
        		}  
        		child { node [draw] {$W_{ij}=0$} 
	        	  edge from parent [->] node [right] {$w_B$}  
        		}
          edge from parent [->] node [right] {$ $}  
        }
    edge from parent [->] node [right] {$\quad1-\psi$} 
    } ;
\end{tikzpicture}

%\includegraphics[scale=0.65]{arbre_Proba.png}
        \caption{\textbf{Tree diagram illustrating the structure of the extended hierarchical model accounting for identification in occupancy}.
        The nodes represent the possible events for the latent occupancy and detection variables, $Z$ and $Y$, respectively associated with the occurrence probabilities $\psi$ and $p$, defined along the branches. The leaves indicate the observed data, $W_{ij}$, recorded during visit $j$ at site $i$, which depend on the detection state $Y_{ij}$ and the associated identification probability : $w_A$ if the species is detected ($Y_{ij}=1$), and $w_B$ otherwise. The detection of the target species ($Y_{ij}=1$) occurs with probability $\psi$ at an occupied site $i$ (i.e $Z_i=1$).}
    \label{fig:tree}
\end{figure}

For each site, the identification record of the target species is compiled on the basis of visits. 
We can derive the probability to observe $w$ ($w=0$ or $1$) at visit $j$ on site $i$ by considering the different possible states for $Y_{ij}$:
\begin{flalign}
    \pi_1(w) & \defeq  Pr\left (W_{ij}  = w\ |Z_i=1 \right)&& \nonumber \\ \label{equ:w1}  
    & = Pr\left (W_{ik}= w,\ Y_{ij}=1\ | Z_i=1\right ) + Pr\left (W_{ij}=  w, Y_{ij}=0\ |Z_i=1\right )&& \nonumber \\
    & = Pr\left (W_{ij}= w\ | Y_{ij}=1\right)  Pr \left ( Y_{ij}=1\ | Z_i=1\right ) + Pr\left (W_{ij}=  w\ | Y_{ik}=0\right ) Pr\left (  Y_{ij}=0\ | Z_i=1\right )&& \\ 
    & = w_A ^{w}(1-w_A)^{1-w} p + (1-w_B)^{w}(w_B)^{1-w} (1-p)&& \nonumber
\end{flalign}
\begin{flalign} 
    \pi_0(w) & \defeq Pr\left (W_{ij}  = w\ |Z_i=0 \right)&& \nonumber \\ \label{equ:w2}
    & = Pr\left (W_{ij}=  w,\ Y_{ij}=0\ |Z_i=0\right )&& \nonumber \\
    & =  Pr\left (W_{ij}=  w\ |  Y_{ij}=0\right ) Pr\left ( Y_{ij}=0\ | Z_i=0\right )&& \\
    & =  (1-w_B)^{w}(w_B)^{1-w}&& \nonumber 
\end{flalign}
\FloatBarrier


For example, at a site visited three times, where the species is identified only during the second visit, the identification history would be $010$. 
Out of these three visits, the occupancy state of the site is unknown but the species was identified once so we combine equations \ref{equ:w1}, \ref{equ:w2}, which account for the site's occupancy state. This may be a true identification; in which case the species is present on the site but not easily identifiable. Otherwise, because this model includes false-positives, the species may have been wrongly identified and the site would not be occupied (Fig. \ref{fig:tree}). Without including false-positives in the identification process, the site would have been necessarily considered occupied.

Conditionally on the site occupancy status and given that the visits are assumed to be independent, the probability to observe the identification history $W_i = (0,1,0)$ is given by:
\begin{align}
    Pr\left ( W_i=(0,1,0) \right) & =
    Pr\left (W_i=(0,1,0),\ Z_i=1\right ) + Pr\left (W_i=(0,1,0),\ Z_i=0\right ) \nonumber \\
    & = Pr\left (W_i=(0,1,0)\ |Z_i=1\right ) Pr\left ( Z_i =1 \right ) %\nonumber  \\
    % & \hspace{2cm}
     + Pr\left (W_i=(0,1,0)\ |Z_i=0\right ) Pr\left ( Z_i =0 \right )  \\ \label{equ:example}
    & = \psi  \pi_1(w_{i1})\pi_1(w_{i2})\pi_1(w_{i3}) + (1-\psi)\pi_0(w_{11})\pi_0(w_{12})\pi_0(w_{13}) \nonumber 
    % \label{equ:example}
\end{align}
Finally, for $S$ independent sites, each  with $J$ independent visits - where $j_i^*$ denotes positive identification - and assuming constant parameters across visits and sites, the model likelihood can be expressed as :

\begin{equation}
  \begin{split}
        L(w_A,w_B,p,\psi\ | data) & = \prod_{i=1}^N Pr(W_i) = \prod_{i=1}^N \left( Pr(W_i, Z_i = 1) +  Pr(W_i, Z_i = 0)  \right) \\ 
        & = \prod_{i=1}^N \Bigg[ \psi \left[ (1-w_B) (1-p) + w_A p \right]^{j_i^*}  \left[ w_B (1-p) + (1-w_A) p \right] ^{J-j_i^*} + \\
        & \qquad\qquad (1-\psi)  w_B^{J-j_i^*}  (1-w_B)^{j_i^*} \Bigg]
    \end{split}
    \label{equ:negloglik}
\end{equation}


\section*{Simulation study}
\subsection*{Classical estimation with a frequentist approach}

In this section, we assess the quality of estimates obtained through maximum likelihood using a simulation study.
Specifically, we aim to assess two key aspects: first, whether incorporating the identification process and accounting for its two types of error leads to more reliable estimates; second, how the number of site visits affects the precision of these estimates.

In order to investigate these points, we carried out simulations by generating 1000 data sets with N=30 sites and J=12 or 36 visits according to our proposed model defined in Equations \eqref{eq:standardmodel}, \eqref{eq:model_w}. The parameter values used to create the matrices of observations were $\psi=0.8$, $p=0.5$, $w_A=0.9$ and $w_B=0.7$. These values were chosen based on a site occupancy study of the Eurasian lynx (Lynx lynx) population in France \parencite{gimenez_tradeoff_2022}.  
After generating the datasets, we applied maximum likelihood estimation by minimizing the negative log-likelihood function to obtain parameter estimates (Equ. \ref{equ:negloglik}). To examine the influence of the number of visits, we compared the precision of estimates between datasets with 12 visits and those with 36 visits.

The results reveal that the occupancy parameter, ${\psi}$, tends to be overestimated when using the original model without the identification. This overestimation occurs because, in the absence of the identification process, all sites with at least one positive identification are assumed to be occupied (mean estimates for 1000 simulations with the original model for 36 visits : $\bar{\hat{\psi}}=1$).

\subsection*{Identifiability issues}

Previous studies have demonstrated that parameter estimates become biased if false-positive detections are not properly accounted for. In particular, the detection probability is underestimated, and occupancy is overestimated \parencite{royle_generalized_2006, mcclintock_unmodeled_2010, miller_improving_2011}.

In our analysis, we used the standard deviation of estimates as a measure of accuracy, which decreases as the number of occasions increases (from $0.22$ for 12 visits to $0.08$ for 36 visits for occupancy probability estimates $\hat{\psi}$)(Fig. \ref{fig:J1236}).
However, despite the increase in available data from 36 visits, the estimates for the detection probability, $\hat{p}$, and the positive identification probability, $\hat{w_A}$, remain biased ($Bias(\hat{p}) = 0.17$ and $Bias(\hat{w_A}) = -0.15$).

One way to address these biases is to fix one of the two parameters, $w_A$ or $p$, then the other can  be estimated without bias (Supplementary \ref{fig:paramfix}).
Such parameter redundancy in the likelihood function is at the core of model identifiability issues (Supplementary \ref{fig:devprof}, \ref{table:devprof})\parencite{gimenez_methods_2004}.

\begin{figure}[h!]
    \centering
    \includegraphics[scale=0.5]{figure1}
    \caption{\textbf{Identifiability issues in Site Occupancy Model accounting for false-positive and false-negative errors in the identification layer}. Histogram and kernel estimates of the distribution of maximum-likelihood estimates for 1000 simulations for J=12 (left column) or J=36 (right column) visits on N=30 sites, and the initial parameter value use to create datasets (in red). Estimates are the occupancy probability $\hat{\psi}$, the detection probability $\hat{p}$, the positive identification probability $\hat{w_A}$ and the negative identification probability $\hat{w_B}$.}
    \label{fig:J1236}
\end{figure}

\subsection*{Addressing identifiability issues with a constraint}

To further address the lack of identifiability in models that incorporate misdetection, \textcite{royle_generalized_2006} suggested to impose constraints on the model. They proposed to set the probability to correctly detect a present species higher than the probability to incorrectly detect it when it is absent. We first explore this recommendation using a frequentist approach, before turning on a Bayesian approach using informative priors in order to solve these identifiability issues.

To adapt the recommended constraint to our model, we apply it on the identification probabilities, such that $w_A > 1 - w_B$. This ensures that the probability of correctly identifying the species is higher than the probability of making a false positive identification.

To evaluate the impact of this constraint, we simulated 1000 datasets with values for the true-positive identification probability $w_A$ and the true-negative identification probability $w_B$ ranging between 0.5 and 0.95. We then estimated the parameters of our site occupancy model accounting for both types of error in the identification layer, using maximum likelihood estimation with and without the constraint.
\FloatBarrier
\begin{figure}[h!]
    \centering
    \includegraphics[scale=0.75]{graphs/figure2.jpeg}
    \caption{\textbf{Bias trend as a function of the probability of correctly identifying the species.} The focus is on parameters likely to be biased by identifiability issues : the detection estimates $\hat{p}$ (on the left), and the correct identification estimates $\hat{w_A}$ (on the right). The bias is contrasted between two optimization cases: under the constraint \textit{(in blue)} stating that the probability of correctly identifying the species is higher than the probability of incorrectly identifying the species, and without the constraint \textit{(in gray)}. The bias is assessed according to the true value of $w_A$ used in the data simulation, and is calculated based on the median and the range between the 0.1 and 0.9 quantiles of the maximum-likelihood estimates.}
    \label{fig:uselesconstrain2}
\end{figure}

The results show that applying the constraint reduces the bias in the detection probability estimates ($\hat{p}$ for values of $w_A$ and $w_B$ around 0.5 ; Supplementary \ref{fig:optcont}). Moreover, regardless of the initial value of $w_A$, the estimates of $\hat{w_A}$ are concentrated around 0.7, which leads to a reduction in bias as the value of  $\hat{w_A}$ (Fig. \ref{fig:J1236}). As $w_A$ and $w_B$ approach higher values, the estimates produced with and without the constraint become more similar.
Nevertheless, while the constraint helps reduce bias, it may not be strong enough to completely eliminate the identifiability issue (Fig. \ref{fig:uselesconstrain2}). This is because, in practice, the true-positive rate, $w_A$, is generally higher than the false-positive rate $1-w_B$ \parencite{guillera-arroita_dealing_2017}.


\subsection*{Using an informative prior to address identifiability issues}

In this section we address the issue of the model identifiability by leveraging knowledge about the risk of misidentifications, even in the absence of additional data sources. We adopt a Bayesian approach, incorporating this knowledge through the use of an informative prior. 

In many situations, it is possible to have a good knowledge of the false-negative rate in the identification process. In particular, we are interested in utilising prior knowledge regarding the sensitivity of the identification process as a means of addressing the redundancy between detection and positive identification parameters, previously described. As the process of species identification is inherently imperfect, its performance is evaluated through the implementation of tests which compare the predicted identifications to the actual outcomes of a verified dataset. Insofar as the underlying truth of the data is not accessible, these performance tests must be carried out beforehand, thus facilitating the acquisition of knowledge regarding the risk of misidentifications. Therefore, the inclusion of additional data sources free of one kind of misidentification is not necessary. 

In the context of sensor data classified by a deep learning algorithm, labelled data are used to evaluate the performance of the classifier before employing it for the classification of unlabeled data \parencite{pichler_machine_2023}. Performance tests are designed to compute metrics that quantify both types of misclassifications. These include the recall defined as the true positive rate (or sensitivity) for each class, and which is of particular interest in the context of identifying one target species \parencite{pichler_machine_2023}. This information is often accessible in the confusion matrix of a classifier, and the transfer learning ensures the consistency of the classifier's performance on other datasets \parencite{tabak_machine_2019,norouzzadeh_deep_2021, velez_evaluation_2023}. Those performance metrics, including sensitivity, may constitute prior knowledge that is more or less informative. Here we examine how the contribution of this external information, integrated into the elicitation of a prior, can be used to address identifiability issues and reduce bias in parameter estimates. We attempt to construct the most suitable prior distribution given the available knowledge about the identification process, and more particularly on the sensitivity of this process modeled by the parameter $w_A$, i.e., the probability that the species will be identified when it is detected.

A highly informative knowledge is characterised by a precise definition of the sensitivity with a median value enhanced by a confidence interval. Consequently, the sensitivity can be expressed as a density distribution with a mean and a standard deviation (e.g. \cite{griffin_modelling_2020} with 0.81 [0.71,0.90] and \cite{tabak_improving_2020} provide the recall values and 95\% confidence intervals for each studied species with \textit{MLWIC2}). In this context, a beta distribution is the most appropriate distribution to elicit a prior on the probability of correctly identifying a species present \parencite{banner_use_2020}. In the case of lesser but still informative knowledge, sensitivity can be defined as a unique value without any confidence interval (e.g \cite{schneider_recognition_2024} give the confusion matrices from their open species recognition models, and the \textcite{Wildlife_Insights_nodate} platform gives its classifier's performance metrics for many species). We then specified a spread beta distribution as a weakly informative prior. In the absence of information concerning the sensitivity of the identification process, it may be reasonably argued that the probability of correctly identifying the target species in an occupancy study is greater than the probability of incorrectly identifying it. This vague knowledge justifies the consideration of a flat uniform distribution ranging from 0.5 to 1 for the positive identification parameter.

Based on \textcite{banner_use_2020} proposition and according to the available knowledge about the sensitivity of the identification process, we study 4 different types of prior for parameter $w_A$ (Supplementary \ref{fig:priorDistribs}) :
\begin{itemize}
    \item a uniform distribution from 0 to 1, as a default non-informative prior for a probability,
    \item a flat uniform distribution ranging from 0.5 to 1, as a vague non-informative prior adapted to the context of identification for occupancy,
    \item a spread-out beta distribution, as a weakly informative prior,    
    \item a tight beta distribution, as a highly informative prior.
\end{itemize}

The beta prior distribution was elicited using a matching method to accurately define its parameters \parencite{denham_geographically_2007,falconer_methods_2022}. Following the approach proposed by \textcite{wu_elicitation_2008} we constructed a unimodal beta distribution through a two-step process. First, we aligned the sensitivity value with the mode of the beta distribution, which represents the most frequent value. Here the sensitivity value is 0.9 according to the values used for the simulations and as a reference to \textcite{gimenez_tradeoff_2022}. Subsequently, we integrated the probability density function by utilizing the confidence interval of the sensitivity as the distribution's range. We simulated 100 observation datasets and we estimated model parameters in a Bayesian framework (using NIMBLE v1.2.0; \cite{valpine_nimble_2024}) for each prior distributions of $w_A$ (the distribution priors of all the others parameters are default prior i.e $\mathcal{U}(0,1)$). We used a block sampler accounting for the correlation between the detection $p$, and the positive identification $w_A$, parameters. The model convergence was analysed for different values of positive identification probability as a simulation parameter (Supplementary \ref{fig:tracePlots}, \ref{fig:SensAnaly}).
\begin{figure}[h!]
    \centering
    \includegraphics[scale=0.5]{figure3}
    \caption{\textbf{Boxplot of the difference between the median values of the posterior distributions and the parameter values calculated from simulated datasets.} Occupancy parameters are set to fixed values to simulate 100 datasets : $\psi=0.8$, $p=0.5$, $w_A=0.9$, $w_B=0.7$. The sensitivity parameter ($w_A$) is introduced as \textbf{(A)} a default non-informative prior with a uniform distribution $\mathcal{U}(0,1)$, \textbf{(B)} a vague prior with a uniform distribution like $\mathcal{U}(0.5,1)$, \textbf{(C)} a weakly informative prior with a beta distribution like $\mathcal{B}(8.8,1.9)$, and \textbf{(D)} a highly informative prior with a beta distribution like $\mathcal{B}(45,5)$.}
    \label{fig:prior_med}
\end{figure}
\FloatBarrier
Using non-informative priors for identification parameters leads to biased posterior distributions, especially for the detection and positive identification parameters. The mean bias associated with the median of the posterior for $\hat{p}$ and $\hat{w_A}$ are $0.13$ and $-0.19$, respectively, when using a default non-informative prior for sensitivity. Notably the negative bias on the positive identification parameter, $w_A$, is not fully compensated by the bias on the detection parameter.
The inference for the detection probability $\hat{p}$ improves when an informative prior for sensitivity is applied. In this case, the mean bias associated with the median of the posterior for $\hat{p}$ decreases to -0.02 with a highly informative prior (Fig. \ref{fig:prior_med}). A vague non-informative prior slightly reduces the mean bias in the median of the posteriors of $\hat{\psi}$. The informative priors used represent two different approaches to integrate information about the identification process, and both perform comparably concerning the estimate of the occupancy probability. Actually, the median values of $\hat{\psi}$ posteriors, obtained for 100 simulations are only weakly affected by the type of prior.


\section*{Discussion}

We proposed a single-species occupancy model that can be applied to various data types, including images, acoustic recordings, and molecular data. This model acknowledges the two-step structure of the observation process, consisting of detection and identification.
Our hierarchical occupancy model considers both detection and identification processes, which are independent sources of errors. On the one hand, we account for false negatives in detection using the detection parameter $p$, and on the other hand, we address identification errors, whether in favor of the target species or not, with parameters $w_A$ and $w_B$.
Initially, we implemented our model within a maximum-likelihood framework, but we encountered biases in some estimates due to model mis-specifications and identifiability issues. By shifting to a Bayesian approach and using informative priors based on identification performance metrics, such as sensitivity, we successfully mitigated these identifiability issues.

The deployment of sensors and molecular techniques generates more data than conventional sampling methods, and because these data are not inherently specific to any species, they require further sorting to identify the target species. Particularly with sensor data, this secondary stage may involve multiple observers, through crowd-sourced projects (e.g. \cite{noauthor_zooniverse_nodate}) for images classification, or expert analysis for acoustic data (e.g. \cite{zwart_use_2014, shonfield_autonomous_2017}). Automated species recognition can reduce processing time, but without human verification which is time-consuming \parencite{barre_accounting_2019, spiers_estimating_2022}, identification errors can distort inferences \parencite{mcclintock_unmodeled_2010, ferguson_occupancy_2015, lonsinger_efficacy_2023}. 
Accounting for these identification errors in addition to detection errors requires developing different versions of the site occupancy model. Firstly, the model developed by \textcite{nichols_multi-scale_2008} considered multiple detection methods at the sampling occasion scale, and so introduced the idea we are following, that a visit on a site may be a set of observations. In essence, dividing a visit into two different detection events is equivalent to the two-stage survey protocol proposed by \textcite{guillera-arroita_dealing_2017}, which we rely on. 
Finally, by reducing data processing time through automation and the absence of human validation, potential identification errors are introduced, which, especially false positives, may have a severe impact on inferences. As the number of model parameters increases to better accommodate different sampling levels, the price to pay is that some parameters become difficult to estimate. Several authors have therefore suggested combining multiple sources of information \parencite{miller_improving_2011, chambert_modeling_2015, guillera-arroita_dealing_2017} to overcome the problem of identifiability. However, since increasing data sources is costly, we propose using performance metrics from the identification process to inform priors. 


In the context of molecular data, a species is detected if its DNA is present in the sample, and it is identified if its DNA is observed in a PCR analysis replicate \parencite{schmidt_site_2013}. Sensitivity is thus defined as the probability of correctly identifying the species, or pathogen, in the replicate. Unlike acoustic or camera trap methods, where detection and identification can be separated, this distinction is more challenging in eDNA surveys, where the sample composition remains unknown until molecular and bioinformatics analysis are performed \parencite{goldberg_critical_2016}. Some studies use additional surveys to verify species presence and calibrate eDNA sensitivity, while others rely on experimental or statistical methods (e.g. \cite{griffin_modelling_2020, mathieu_systematic_2020}). The use of positive control involving foreign DNA, can help to identify PCR inhibition and provide information on the false-positive rate (e.g. \cite{goldberg_critical_2016, furlan_framework_2016})\parencite{hyatt_diagnostic_2007}. Nevertheless, quantifying sensitivity remains challenging across studies using similar methodologies due to high variability in taxa, environmental, and experimental conditions \parencite{thomsen_monitoring_2012, keller_tracking_2022, gold_comparison_2023}. Despite this, eDNA is generally more sensitive than other sampling methods \parencite{darling_molecules_2011}, though this heightened sensitivity may increase the likelihood of false positives \parencite{cristescu_uses_2018}. Taking into account the identification process is therefore crucial, although the positive identification rate ($w_A$) must be close enough to 1 to guarantee the convergence of the model.

The main limitation of our approach lies in the fact that we need to gather knowledge on the performance of the identification process to construct a relevant informative prior. While this knowledge is necessary, it is still less costly than incorporating additional data sources, especially if sensitivity information is provided by another study, or as a parameter of the identification tool (e.g. \cite{tabak_improving_2020}, \cite{rigoudy_deepfaune_2023}).
Indeed, we suggest that when using deep learning algorithms for species classification, or following a molecular and bioinformatics pipeline for eDNA, the performance metrics of the methods should be made accessible.
Simulations indicate that even with non-informative priors, our model produces reliable posterior estimates of the presence parameter ($\psi$). When only presence is of interest, we recommend using this model with non-informative priors to handle misidentifications and detection errors while disregarding identifiability issues in the detection parameter. However, when the detection parameter is of concern, using an informative prior is necessary to address parameter redundancy.
\textcite{cruickshank_quantifying_2019} successfully avoided identifiability issues related to false-positive errors by integrating informative prior based reasonable assumptions from volunteer-collected monitoring data. Similarly, our approach, which incorporates prior information about the identification process, produces robust posterior estimates and provides an alternative to approaches requiring additional datasets.
Also, as in many studies using a Bayesian approach, the choice of a wrong prior for a parameter may cause bias in the definition of the posterior distribution for this parameter \parencite{northrup_comment_2018}.


Passive sensors like camera traps and autonomous recording units offer valuable opportunities for addressing a wide range of ecological and conservation questions.  Combined with approaches like eDNA sampling, these technologies enable ecologists to collect data at large spatial scales or fine temporal resolutions and study cryptic species \parencite{ross_passive_2023, sahu_environmental_2023}. For such large and complex datasets, accurate taxonomic identification is challenging, but accounting for the noise generated during processing is essential. In this context, our proposed model can be included in the ecologist’s toolbox for analyzing sensor and molecular biological data to address questions in conservation biology, wildlife management and disease ecology.


\newpage
\counterwithin{figure}{section}
\counterwithin{table}{section}
\appendix
\section{Supplementary Results}\label{sR}
\subsection{Identifiability issues}
\begin{figure}[h!]
    \centering
    \includegraphics[scale=0.5]{figureA1}
    \caption{\textbf{Distribution of maximum-likelihood estimates for 1000 simulations} when a parameter is set to a constant value (in columns).Detection ($p$) and identification parameters ($w_A$ and $w_B$) are successively excluded from the estimation, since their value are fixed in the expression of the likelihood function.}
    \label{fig:paramfix}
\end{figure}

$\hat{w_A}$ or $\hat{p}$ are estimated without bias when the other parameter is set to a fixed value in the expression of likelihood. This result reflects parameter redundancy in the likelihood function.

We consider the profile deviance on $p$ to investigate model identifiability. 

\begin{figure}[h!]
    \centering
    \includegraphics[scale=0.75]{figureA2}
    \caption{Profile deviance on $p$}
    \label{fig:devprof}
\end{figure}

Deviance ($-2Log-Likelihood$) is constant for $p$ greater than 0.45, beyond this value the model is not identifiable, which means that $\hat{p}$ and $\hat{w_A}$ cannot be distinguished.

The model is not globally identifiable \parencite{cole_determining_2010} since there are different sets of parameters that give rise to the same likelihood function value.

\begin{table}[h!]
    \centering
    \caption{Profile deviance on detection parameter $p$}
    \begin{tabular}{c c c c c}
    \hline
    $\hat{\psi}$ & $p$ & $\hat{w_A}$ & $\hat{w_B}$ & \textbf{Likelihood} \\
    \hline
    0.7419 & 0.9 & 0.6592 & 0.6777 & \textbf{719.6974} \\
    0.7421 & 0.8 & 0.7013 & 0.6777 & \textbf{719.6974} \\
    0.7419 & 0.7 & 0.7554 & 0.6778 & \textbf{719.6974} \\
    0.7422 & 0.6 & 0.8276 & 0.6776 & \textbf{719.6974} \\
    0.7419 & 0.5 & 0.9286 & 0.6777 & \textbf{719.6974} \\
    % 0.7325 & 0.4 & 1 & 0.6470 & \textbf{720.5211} \\
    \end{tabular}
    \label{table:devprof}
\end{table}

As pointed out by \textcite{royle_generalized_2006}, including false positives raises concerns about model identifiability . To address this issue of parameter redundancy, the authors proposed to set a constraint during likelihood optimization. Specifically, they suggest ensuring that the probability of correctly detecting a species is higher than the probability of falsely detecting it. Applying this constraint to our model with an identification layer means that correctly identifying the target species is more likely than falsely identifying it when it has not been detected. 

\begin{figure}[h!]
    \centering
    \includegraphics[scale=0.5]{graphs/figureA3.png}
    \caption{\textbf{Distribution of $\hat{\psi}$ and $\hat{p}$ for 1000 simulated data sets for different values of identification parameters in the simulated data.} With $w_A$ set between 0.5 and 0.95 (top) and $w_B$ set between 0.5 and 0.95 (bottom). Distributions of occupancy ($\hat{\psi}$) and detection $\hat{p}$ parameters are the results of optimization under the constraint $\hat{w_A} > 1 - \hat{w_B}$ (in gray) and without it (in blue). The true value of parameters are indicated by the red vertical bar.}
    \label{fig:optcont}
\end{figure}

The constraint proposed does not help to fix the estimation issue in the detection probability, however for small values of $w_A$ or $p$, close to 0.5, occupancy estimates are reliable.

\subsection{Using an informative prior to address identifiability issues}
We evaluate the posterior distributions of the occupancy estimates according to four priors with different level of informativeness for the positive identification parameter, $w_A$, called sensitivity.

\begin{figure}[h!]
    \centering
    \includegraphics[scale=0.75]{cedram/graphs/figureA4.png}
    \caption{\textbf{Prior distributions for the positive identification parameter or sensitivity $w_A$.} Non informative prior (in blue) are uniform distributions : from 0 to 1 (in dark blue) and from 0.5 to 1 (in light blue). Informative priors (in orange) are beta distributions such as $\mathcal{B}(8.8,1.9)$ is weakly informative (in light orange) and $\mathcal{B}(76,9.3)$ is highly informative (in dark orange).}
    \label{fig:priorDistribs}
\end{figure}
We elicited the beta priors by solving a 2 equations system explicating the mode and the density probability function with the beta distribution parameters,  $\alpha$ and $\beta$, unknown (in the manner of the location and intervals method of \textcite{wu_elicitation_2008}) :
\begin{gather}
    mode = \frac{\alpha-1}{\alpha + \beta -2}\\
    \int_0^R \frac{x^{\alpha-1}(1-x)^{\beta-1}}{\mathcal{B}(\alpha,\beta)}dx - 0.01=0 \text{ for }P(x<R)=0.01
\end{gather}
For both priors the mode is set to 0.9 which is the value chosen to simulate data. $R$ is defined as the threshold below which the probability to find the value of sensitivity is nearly null : it is 0.5 in the case of a weakly informative prior and 0.8 in the case of the highly informative one.

We ran with NIMBLE (v1.2.0; \cite{valpine_nimble_2024}) 2 chains on 4000 iterations following a 1000 iterations burn-in period. We assessed the model convergence through the R-hat and the trace and density plots (MCMCvis R package v0.16.3; \cite{youngflesh_mcmcvis_2023}), for each alternative priors .

\begin{figure}[h!]
    \centering
    \includegraphics[scale=0.4]{cedram/graphs/figureA5.png}
    \caption{\textbf{Chain trace and density plots of occupancy, $\hat{\psi}$, and detection, $\hat{p}$, posterior distribution, according 4 different priors on sensitivity parameter, $w_A$.} On each of the 4 panels, the trace plots (on the left) represent the evolution of both chains on 4000 iterations, and the density plots (on the right) represent the posterior distribution for each chain. 
    The distribution priors on $w_A$ are (\textbf{A}) $\mathcal{U}(0,1)$,  (\textbf{B}) $\mathcal{U}(0.5,1)$, (\textbf{C}) $\mathcal{B}(8.8,1.9)$ and (\textbf{D}) $\mathcal{B}(76,9.3)$.}
    \label{fig:tracePlots}
\end{figure}
\FloatBarrier
Chains convergence is reached for $\psi$ whatever the prior on $w_A$, however only the most informative prior enable a satisfying mix of chains for the detection parameter $p$ (R-hat=1.01).

Finally, we drove a sensitivity analysis for 3 values of $w_A$ (0.2, 0.5 and 0.8) used to simulate data. We used a highly informative prior in order to evaluate the impact of the value of $w_A$ on the convergence. 
\begin{figure}[h!]
    \centering
    \includegraphics[scale=0.5]{cedram/graphs/figureA6.png}
    \caption{\textbf{Sensitivity analysis of the extended occupancy model using an highly informative on the positive identification parameter, $w_A$.} Data are simulated for 30 sites visited 36 times with fixed generative values (red dashed line) except for $w_A$.}
    \label{fig:SensAnaly}
\end{figure}
The chains for the occupancy estimates do not converge when the positive identification rate is below 0.5, though this scenario seems unrealistic. This model should only be used when the sensitivity of the identification process is high (greater than 0.75). Indeed, if sensitivity falls below this threshold, the identification process should be considered too underperforming for use in occupancy studies. 

\FloatBarrier
\section*{Acknowledgements}
We would like to acknowledge the assistance of ChatGPT, a language model developed by OpenAI, in improving the clarity and quality of the writing in this manuscript.


\section*{Fundings}

This project has received financial support from the CNRS through the MITI interdisciplinary programs. 

\section*{Conflict of interest disclosure}

The authors declare that they comply with the PCI rule of having no financial conflicts of interest in relation to the content of the article.

\section*{Data, script, code, and supplementary information availability}

Script and codes are available online (\url{https://zenodo.org/doi/10.5281/zenodo.11121903}; Monchy et al., 2024)\\


%\nocite[{*}]
\printbibliography 

\end{document}
