## Supplementary material for "Using informative priors to account for identifiability issues in occupancy models with identification errors": latex files: references2.bib

@article{zwart_use_2014,
	title = {The {Use} of {Automated} {Bioacoustic} {Recorders} to {Replace} {Human} {Wildlife} {Surveys}: {An} {Example} {Using} {Nightjars}},
	volume = {9},
	issn = {1932-6203},
	shorttitle = {The Use of Automated Bioacoustic Recorders to Replace Human Wildlife Surveys},
	url = {https://dx.plos.org/10.1371/journal.pone.0102770},
	doi = {10.1371/journal.pone.0102770},
	language = {en},
	number = {7},
	urldate = {2024-02-02},
	journal = {PLoS ONE},
	author = {Zwart, Mieke C. and Baker, Andrew and McGowan, Philip J. K. and Whittingham, Mark J.},
	month = jul,
	year = {2014},
	pages = {e102770},
}


@online{noauthor_zooniverse_nodate,
	title = {Zooniverse},
	url = {https://www.zooniverse.org/},
	abstract = {The Zooniverse is the world's largest and most popular platform for people-powered research.},
	language = {en},
	urldate = {2024-09-20},
}


@online{Wildlife_Insights_nodate,
	title = {Wildlife Insights},
    abstract={The Wildlife Insights platform uses AI technology to drastically reduce how long it takes to convert camera trap data into usable biodiversity insights. Anyone who collects camera trap data can use Wildlife Insights to upload data to Google Cloud and access AI models trained to automatically classify camera trap images.},
	url = {https://www.wildlifeinsights.org/about-wildlife-insights-ai},
	urldate = {2024-09-23},
}


@article{schneider_recognition_2024,
	title = {Recognition of {European} mammals and birds in camera trap images using deep neural networks},
	copyright = {© 2024 The Author(s). IET Computer Vision published by John Wiley \& Sons Ltd on behalf of The Institution of Engineering and Technology.},
	issn = {1751-9640},
	url = {https://onlinelibrary.wiley.com/doi/abs/10.1049/cvi2.12294},
	doi = {10.1049/cvi2.12294},
	language = {en},
	urldate = {2024-09-19},
	journal = {IET Computer Vision},
	author = {Schneider, Daniel and Lindner, Kim and Vogelbacher, Markus and Bellafkir, Hicham and Farwig, Nina and Freisleben, Bernd},
	year = {2024},
	keywords = {computer vision, convolutional neural nets, image classification, image recognition, neural nets},
}


@article{rigoudy_deepfaune_2023,
	title = {The {DeepFaune} initiative: a collaborative effort towards the automatic identification of {European} fauna in camera trap images},
	volume = {69},
	issn = {1439-0574},
	shorttitle = {The {DeepFaune} initiative},
	url = {https://doi.org/10.1007/s10344-023-01742-7},
	doi = {10.1007/s10344-023-01742-7},
	language = {en},
	number = {6},
	urldate = {2024-09-19},
	journal = {European Journal of Wildlife Research},
	author = {Rigoudy, Noa and Dussert, Gaspard and Benyoub, Abdelbaki and Besnard, Aurélien and Birck, Carole and Boyer, Jérome and Bollet, Yoann and Bunz, Yoann and Caussimont, Gérard and Chetouane, Elias and Carriburu, Jules Chiffard and Cornette, Pierre and Delestrade, Anne and De Backer, Nina and Dispan, Lucie and Le Barh, Maden and Duhayer, Jeanne and Elder, Jean-François and Fanjul, Jean-Baptiste and Fonderflick, Jocelyn and Froustey, Nicolas and Garel, Mathieu and Gaudry, William and Gérard, Agathe and Gimenez, Olivier and Hemery, Arzhela and Hemon, Audrey and Jullien, Jean-Michel and Knitter, Daniel and Malafosse, Isabelle and Marginean, Mircea and Ménard, Louise and Ouvrier, Alice and Pariset, Gwennaelle and Prunet, Vincent and Rabault, Julien and Randon, Malory and Raulet, Yann and Régnier, Antoine and Ribière, Romain and Ricci, Jean-Claude and Ruette, Sandrine and Schneylin, Yann and Sentilles, Jérôme and Siefert, Nathalie and Smith, Bethany and Terpereau, Guillaume and Touchet, Pierrick and Thuiller, Wilfried and Uzal, Antonio and Vautrain, Valentin and Vimal, Ruppert and Weber, Julian and Spataro, Bruno and Miele, Vincent and Chamaillé-Jammes, Simon},
	month = oct,
	year = {2023},
	keywords = {Artificial Intelligence, Camera trap, Classification, Deep learning, Europe, France, Software},
	pages = {113},
}


@article{pichler_machine_2023,
	title = {Machine learning and deep learning—{A} review for ecologists},
	volume = {14},
	copyright = {© 2023 The Authors. Methods in Ecology and Evolution published by John Wiley \& Sons Ltd on behalf of British Ecological Society.},
	issn = {2041-210X},
	url = {https://onlinelibrary.wiley.com/doi/abs/10.1111/2041-210X.14061},
	doi = {10.1111/2041-210X.14061},
	language = {en},
	number = {4},
	urldate = {2024-09-18},
	journal = {Methods in Ecology and Evolution},
	author = {Pichler, Maximilian and Hartig, Florian},
	year = {2023},
	keywords = {artificial intelligence, big data, causal inference, deep learning, machine learning},
	pages = {994--1016},
}


@Manual{valpine_nimble_2024,
    title = {{NIMBLE}: {MCMC}, Particle Filtering, and Programmable Hierarchical Modeling},
    author = {Perry {de Valpine} and Christopher Paciorek and Daniel Turek and Nick Michaud and Cliff Anderson-Bergman and Fritz Obermeyer and Claudia {Wehrhahn Cortes} and Abel Rodrìguez and Duncan {Temple Lang} and Sally Paganin},
    url = {https://cran.r-project.org/package=nimble},
    year = {2024},
    version = {1.2.0},
    note = {{R} package version 1.2.0},
    doi = {10.5281/zenodo.1211190},
}


 @article{youngflesh_mcmcvis_2023,
    title = {MCMCvis: Tools to visualize, manipulate, and summarize MCMC output},
    author = {Casey Youngflesh},
    year = {2018},
    journal = {Journal of Open Source Software},
    volume = {3},
    number = {24},
    doi = {10.21105/joss.00640},
    pages = {640},
}

@article{cristescu_uses_2018,
	title = {Uses and {Misuses} of {Environmental} {DNA} in {Biodiversity} {Science} and {Conservation}},
	volume = {49},
	issn = {1543-592X, 1545-2069},
	url = {https://www.annualreviews.org/content/journals/10.1146/annurev-ecolsys-110617-062306},
	doi = {10.1146/annurev-ecolsys-110617-062306},
	language = {fr},
	number = {Volume 49, 2018},
	urldate = {2024-08-30},
	journal = {Annual Review of Ecology, Evolution, and Systematics},
	author = {Cristescu, Melania E. and Hebert, Paul D. N.},
	month = nov,
	year = {2018},
	pages = {209--230},
}

@article{hartig_novel_2024,
	title = {Novel community data in ecology-properties and prospects},
	volume = {39},
	issn = {0169-5347},
	url = {https://www.cell.com/trends/ecology-evolution/abstract/S0169-5347(23)00265-3},
	doi = {10.1016/j.tree.2023.09.017},
	language = {English},
	number = {3},
	urldate = {2024-08-30},
	journal = {Trends in Ecology \& Evolution},
	author = {Hartig, Florian and Abrego, Nerea and Bush, Alex and Chase, Jonathan M. and Guillera-Arroita, Gurutzeta and Leibold, Mathew A. and Ovaskainen, Otso and Pellissier, Loïc and Pichler, Maximilian and Poggiato, Giovanni and Pollock, Laura and Si-Moussi, Sara and Thuiller, Wilfried and Viana, Duarte S. and Warton, David I. and Zurell, Damaris and Yu, Douglas W.},
	month = mar,
	year = {2024},
	pmid = {37949795},
	keywords = {camera traps, community ecology, environmental DNA (eDNA), joint species distribution models (jSDMs), macroecology, passive acoustic monitoring},
	pages = {280--293},
}


@misc{kirchhof_uncertainties_2024,
	title = {Uncertainties of {Latent} {Representations} in {Computer} {Vision}},
	url = {http://arxiv.org/abs/2408.14281},
	doi = {10.15496/publikation-98103},
	language = {en},
	urldate = {2024-08-27},
	author = {Kirchhof, Michael},
	month = aug,
	year = {2024},
	keywords = {Computer Science - Artificial Intelligence, Computer Science - Computer Vision and Pattern Recognition, Computer Science - Machine Learning},
}

@article{wu_elicitation_2008,
	title = {Elicitation of a {Beta} {Prior} for {Bayesian} {Inference} in {Clinical} {Trials}},
	volume = {50},
	issn = {1521-4036},
	url = {https://onlinelibrary.wiley.com/doi/abs/10.1002/bimj.200710390},
	doi = {10.1002/bimj.200710390},
	language = {en},
	number = {2},
	urldate = {2024-08-14},
	journal = {Biometrical Journal},
	author = {Wu, Yujun and Shih, Weichung J. and Moore, Dirk F.},
	year = {2008},
	keywords = {Bayesian inference, Beta distribution, Clinical trials, Imprecise assessment, Prior and posterior distribution},
	pages = {212--223},
}

@article{hyatt_diagnostic_2007,
	title = {Diagnostic assays and sampling protocols for the detection of {Batrachochytrium} dendrobatidis},
	volume = {73},
	issn = {0177-5103, 1616-1580},
	url = {http://www.int-res.com/abstracts/dao/v73/n3/p175-192/},
	doi = {10.3354/dao073175},
	language = {en},
	urldate = {2024-07-25},
	journal = {Diseases of Aquatic Organisms},
	author = {Hyatt, Ad and Boyle, Dg and Olsen, V and Boyle, Db and Berger, L and Obendorf, D and Dalton, A and Kriger, K and Hero, M and Hines, H and Phillott, R and Campbell, R and Marantelli, G and Gleason, F and Colling, A},
	month = jan,
	year = {2007},
	pages = {175--192},
}


@article{darling_molecules_2011,
	title = {From molecules to management: {Adopting} {DNA}-based methods for monitoring biological invasions in aquatic environments},
	volume = {111},
	issn = {0013-9351},
	shorttitle = {From molecules to management},
	url = {https://www.sciencedirect.com/science/article/pii/S0013935111000569},
	doi = {10.1016/j.envres.2011.02.001},
	number = {7},
	urldate = {2024-07-24},
	journal = {Environmental Research},
	author = {Darling, John A. and Mahon, Andrew R.},
	month = oct,
	year = {2011},
	keywords = {Aquatic invasive species, DNA-based monitoring, Detection, Management, Uncertainty},
	pages = {978--988},
}

@article{mathieu_systematic_2020,
	title = {A {Systematic} {Review} of {Sources} of {Variability} and {Uncertainty} in {eDNA} {Data} for {Environmental} {Monitoring}},
	volume = {8},
	issn = {2296-701X},
	url = {https://www.frontiersin.org/journals/ecology-and-evolution/articles/10.3389/fevo.2020.00135/full},
	doi = {10.3389/fevo.2020.00135},
	language = {English},
	urldate = {2024-07-24},
	journal = {Frontiers in Ecology and Evolution},
	author = {Mathieu, Chloé and Hermans, Syrie M. and Lear, Gavin and Buckley, Thomas R. and Lee, Kevin C. and Buckley, Hannah L.},
	month = may,
	year = {2020},
	keywords = {Spatiotemporal scale, bioinformatics, eDNA, experimental design, uncertainty, variability},
}


@article{furlan_framework_2016,
	title = {A framework for estimating the sensitivity of {eDNA} surveys},
	volume = {16},
	copyright = {© 2015 John Wiley \& Sons Ltd},
	issn = {1755-0998},
	url = {https://onlinelibrary.wiley.com/doi/abs/10.1111/1755-0998.12483},
	doi = {10.1111/1755-0998.12483},
	language = {en},
	number = {3},
	urldate = {2024-07-24},
	journal = {Molecular Ecology Resources},
	author = {Furlan, Elise M. and Gleeson, Dianne and Hardy, Christopher M. and Duncan, Richard P.},
	year = {2016},
	keywords = {detection probability, detection sensitivity, distribution, eDNA, environmental DNA, species-specific detection},
	pages = {641--654},
}


@article{goldberg_critical_2016,
	title = {Critical considerations for the application of environmental {DNA} methods to detect aquatic species},
	volume = {7},
	copyright = {© 2016 The Authors. Methods in Ecology and Evolution published by John Wiley \& Sons Ltd on behalf of the British Ecological Society},
	issn = {2041-210X},
	url = {https://onlinelibrary.wiley.com/doi/abs/10.1111/2041-210X.12595},
	doi = {10.1111/2041-210X.12595},
	language = {en},
	number = {11},
	urldate = {2024-07-24},
	journal = {Methods in Ecology and Evolution},
	author = {Goldberg, Caren S. and Turner, Cameron R. and Deiner, Kristy and Klymus, Katy E. and Thomsen, Philip Francis and Murphy, Melanie A. and Spear, Stephen F. and McKee, Anna and Oyler-McCance, Sara J. and Cornman, Robert Scott and Laramie, Matthew B. and Mahon, Andrew R. and Lance, Richard F. and Pilliod, David S. and Strickler, Katherine M. and Waits, Lisette P. and Fremier, Alexander K. and Takahara, Teruhiko and Herder, Jelger E. and Taberlet, Pierre},
	year = {2016},
	keywords = {biodiversity, eDNA, invasive species, non-destructive sampling, quantitative PCR, reporting guidelines},
	pages = {1299--1307},
}

@article{keller_tracking_2022,
	title = {Tracking an invasion front with environmental {DNA}},
	volume = {32},
	copyright = {© 2022 The Authors. Ecological Applications published by Wiley Periodicals LLC on behalf of The Ecological Society of America.},
	issn = {1939-5582},
	url = {https://onlinelibrary.wiley.com/doi/abs/10.1002/eap.2561},
	doi = {10.1002/eap.2561},
	language = {en},
	number = {4},
	urldate = {2024-07-24},
	journal = {Ecological Applications},
	author = {Keller, Abigail G. and Grason, Emily W. and McDonald, P. Sean and Ramón-Laca, Ana and Kelly, Ryan P.},
	year = {2022},
	keywords = {Bayesian modeling, Carcinus maenas, European green crab, N-mixture modeling, environmental DNA, false-positive probability, invasion front, invasive species management},
	pages = {e2561},
}


@article{banner_use_2020,
	title = {The use of {Bayesian} priors in {Ecology}: {The} good, the bad and the not great},
	volume = {11},
	issn = {2041-210X},
	shorttitle = {The use of {Bayesian} priors in {Ecology}},
	url = {https://onlinelibrary.wiley.com/doi/abs/10.1111/2041-210X.13407},
	doi = {10.1111/2041-210X.13407},
	language = {en},
	number = {8},
	urldate = {2024-07-23},
	journal = {Methods in Ecology and Evolution},
	author = {Banner, Katharine M. and Irvine, Kathryn M. and Rodhouse, Thomas J.},
	year = {2020},
	keywords = {Bayesian hierarchical models, good statistical practice, sensitivity analysis, subjective priors},
	pages = {882--889},
}


@article{goldman_integrating_2023,
	title = {Integrating multiple sign types to improve occupancy estimation for inconspicuous species},
	volume = {13},
	issn = {2045-7758},
	url = {https://onlinelibrary.wiley.com/doi/10.1002/ece3.10019},
	doi = {10.1002/ece3.10019},
	abstract = {Abstract Standard occupancy models enable unbiased estimation of occupancy by accounting for observation errors such as missed detections (false negatives) and, less commonly, incorrect detections (false positives). Occupancy models are fitted to data from repeated site visits in which surveyors record evidence of species presence. Use of indirect sign (e.g., scat, tracks) as evidence of presence can vastly improve survey efficiency for inconspicuous species but can also introduce additional sources of error. We developed a ?multi-sign? occupancy approach to model the detection process separately for unique sign types and used this method to improve estimates of occupancy dynamics for an inconspicuous species, the American pika (Ochotona princeps). We investigated how estimates of pika occupancy and environmental drivers differed under four increasingly realistic representations of the observation process: (1) perfect detection (commonly assumed for modeling pika occupancy), (2) standard occupancy model (single observation process without possibility of false detection), (3) multi-sign with no false detections (non-false positive model), and (4) multi-sign with false detections (full model). For the multi-sign occupancy models, we modeled the detection of each sign type (fresh scat, fresh haypiles, pika calls, and pika sightings) separately as a function of climatic and environmental covariates. Estimates of occupancy processes and inferences about environmental drivers were sensitive to different detection models. Simplified representations of the detection processes generally resulted in higher occupancy estimates and higher turnover rates than the full multi-sign model. Environmental drivers also varied in their influence on occupancy models, where (e.g.) forb cover was estimated to more strongly influence occupancy in the full multi-sign model than the simpler models. As has been reported previously in other contexts, unmodeled heterogeneity in the observation process can lead to biases in occupancy processes and uncertainty in the relationships between occupancy and environmental covariates. Overall, our multi-sign approach to dynamic occupancy modeling, which accounts for spatio-temporal variation in reliability among sign types, has strong potential to generate more realistic estimates of occupancy dynamics for inconspicuous species.},
	number = {5},
	urldate = {2024-06-28},
	journal = {Ecology and Evolution},
	author = {Goldman, Mia R. and Shinderman, Matthew and Jeffress, Mackenzie R. and Rodhouse, Thomas J. and Shoemaker, Kevin T.},
	month = may,
	year = {2023},
	keywords = {American pika, detection probability, false negative, false positive, indirect sign, multi-sign, occupancy model, population dynamics},
	pages = {e10019},
}


@article{denham_geographically_2007,
	title = {Geographically {Assisted} {Elicitation} of {Expert} {Opinion} for {Regression} {Models}},
	volume = {2},
	abstract = {One of the perceived strengths of Bayesian modelling is the ability to include prior information. Although objective or noninformative priors might be preferred in some situations, in many other applications the Bayesian framework oﬀers a real opportunity to formally combine data with information available from experts. The question addressed in this paper is how to elicit this information in a form suitable for prior modelling. Particular attention is paid to geographic data for which maps might be used to assist in the elicitation. Two case studies are used to illustrate the methodology: estimation of city house prices and prediction of presence of a rare species.},
	language = {en},
	number = {1},
	journal = {Bayesian Analysis},
	author = {Denham, Robert and Mengersen, Kerrie},
	year = {2007},
	pages = {99--136},
}


@article{mccarthy_profiting_2005,
	title = {Profiting from prior information in {Bayesian} analyses of ecological data},
	volume = {42},
	issn = {1365-2664},
	url = {https://onlinelibrary.wiley.com/doi/abs/10.1111/j.1365-2664.2005.01101.x},
	doi = {10.1111/j.1365-2664.2005.01101.x},
	abstract = {1 Most ecological studies include prior information only implicitly, usually in their design or the discussion of results. In this study, two examples demonstrate that using Bayesian statistics to incorporate basic ecological principles and prior data can be very cost-effective for increasing confidence in ecological research. 2 The first example is based on examining the effects of an experimental manipulation of the habitat of mulgara Dasycercus cristicauda, a marsupial of inland Australia. The second example is based on observational mark–recapture data to estimate the annual survival of the European dipper Cinclus cinclus, a passerine in France. 3 In the mulgara example, the prior information obtained from an observational study increased confidence that there was an adverse effect of experimental habitat manipulation on the species. The results suggested that the capture rate of mulgara was reduced to approximately one-quarter by reduction of vegetation cover. 4 In the European dipper example, prior information based on the body mass of the species and estimates of annual survival of other European passerines was shown to be worth between 1 and 5 years of mark–recapture field data. 5 Synthesis and applications. Body mass can be used to predict annual survival of European passerines and other animals. Results of observational studies can provide prior information in experimental studies of impacts of habitat change. By using Bayesian methods, such prior information, if represented in a coherent and logical way, can be cost-effective for adding certainty to ecological studies.},
	language = {en},
	number = {6},
	urldate = {2024-03-29},
	journal = {Journal of Applied Ecology},
	author = {McCarthy, Michael A. and Masters, Pip},
	year = {2005},
	keywords = {Cinclus cinclus, Dasycercus cristicauda, Markov chain Monte Carlo, habitat manipulation, mark–recapture, survival},
	pages = {1012--1019},
}


@article{falconer_methods_2022,
	title = {Methods for Eliciting Informative Prior Distributions: A Critical Review},
	volume = {19},
	issn = {1545-8490},
	url = {https://pubsonline.informs.org/doi/10.1287/deca.2022.0451},
	doi = {10.1287/deca.2022.0451},
	shorttitle = {Methods for Eliciting Informative Prior Distributions},
	pages = {189--204},
	number = {3},
	journaltitle = {Decision Analysis},
	author = {Falconer, Julia R. and Frank, Eibe and Polaschek, Devon L. L. and Joshi, Chaitanya},
	urldate = {2024-09-25},
	date = {2022-09},
	keywords = {Bayesian inference, informative priors, prior distribution, prior elicitation, uncertainty},
}


@article{choy_elicitation_2009,
	title = {Elicitation by design in ecology: using expert opinion to inform priors for {Bayesian} statistical models},
	volume = {90},
	copyright = {© 2009 by the Ecological Society of America},
	issn = {1939-9170},
	shorttitle = {Elicitation by design in ecology},
	url = {https://onlinelibrary.wiley.com/doi/abs/10.1890/07-1886.1},
	doi = {10.1890/07-1886.1},
	abstract = {Bayesian statistical modeling has several benefits within an ecological context. In particular, when observed data are limited in sample size or representativeness, then the Bayesian framework provides a mechanism to combine observed data with other “prior” information. Prior information may be obtained from earlier studies, or in their absence, from expert knowledge. This use of the Bayesian framework reflects the scientific “learning cycle,” where prior or initial estimates are updated when new data become available. In this paper we outline a framework for statistical design of expert elicitation processes for quantifying such expert knowledge, in a form suitable for input as prior information into Bayesian models. We identify six key elements: determining the purpose and motivation for using prior information; specifying the relevant expert knowledge available; formulating the statistical model; designing effective and efficient numerical encoding; managing uncertainty; and designing a practical elicitation protocol. We demonstrate this framework applies to a variety of situations, with two examples from the ecological literature and three from our experience. Analysis of these examples reveals several recurring important issues affecting practical design of elicitation in ecological problems.},
	language = {en},
	number = {1},
	urldate = {2024-03-29},
	journal = {Ecology},
	author = {Choy, Samantha Low and O'Leary, Rebecca and Mengersen, Kerrie},
	year = {2009},
	keywords = {design, ecology, expert elicitation, framework, informative Bayesian analysis, prior information, protocol},
	pages = {265--277},
}

@article{cruickshank_quantifying_2019,
	title = {Quantifying data quality in a citizen science monitoring program: {False} negatives, false positives and occupancy trends},
	volume = {1},
	copyright = {© 2019 The Authors. Conservation Science and Practice published by Wiley Periodicals, Inc. on behalf of Society for Conservation Biology},
	issn = {2578-4854},
	shorttitle = {Quantifying data quality in a citizen science monitoring program},
	url = {https://onlinelibrary.wiley.com/doi/abs/10.1111/csp2.54},
	doi = {10.1111/csp2.54},
	abstract = {Data collected by volunteers are an important source of information used in species management decisions, yet concerns are often raised over the quality of such data. Two major forms of error exist in occupancy datasets; failing to observe a species when present (imperfect detection—also known as false negatives), and falsely reporting a species as present (false-positive errors). Estimating these rates allows us to quantify volunteer data quality, and may prevent the inference of erroneous trends. We use a new parameterization of a dynamic occupancy model to estimate and adjust for false-negative and false-positive errors, producing accurate estimates of occupancy. We validated this model using simulations and applied it to 12 species datasets collected from a 15-year, large-scale volunteer amphibian monitoring program. False-positive rates were low for most, but not all, species, and accounting for these errors led to quantitative differences in occupancy, although trends remained consistent even when these effects were ignored. We present a model that represents an intuitive way of quantifying the quality of volunteer monitoring datasets, and which can produce unbiased estimates of occupancy despite the presence of multiple types of observation error. Importantly, this allows the quality of volunteer monitoring data to be assessed without relying on comparisons with expert data.},
	language = {en},
	number = {7},
	urldate = {2024-03-29},
	journal = {Conservation Science and Practice},
	author = {Cruickshank, Sam S. and Bühler, Christoph and Schmidt, Benedikt R.},
	year = {2019},
	keywords = {citizen science, false-positive, imperfect detection, observation bias, occupancy modeling, trend, volunteer monitoring},
	pages = {e54},
}

@article{northrup_comment_2018,
	title = {A comment on priors for {Bayesian} occupancy models},
	volume = {13},
	issn = {1932-6203},
	url = {https://journals.plos.org/plosone/article?id=10.1371/journal.pone.0192819},
	doi = {10.1371/journal.pone.0192819},
	language = {en},
	number = {2},
	urldate = {2023-09-01},
	journal = {PLOS ONE},
	author = {Northrup, Joseph M. and Gerber, Brian D.},
	year = {2018},
    publisher = {Public Library of Science},
	keywords = {Bayesian method, Distribution curves, Ecological niches, Endangered species, Normal distribution, Probability distribution, Simulation and modeling, Theoretical ecology},
	pages = {1-13},
}

@article{morris_neglected_2015,
	title = {The neglected tool in the {Bayesian} ecologist's shed: a case study testing informative priors' effect on model accuracy},
	volume = {5},
	copyright = {© 2014 The Authors. Ecology and Evolution published by John Wiley \& Sons Ltd.},
	issn = {2045-7758},
	shorttitle = {The neglected tool in the {Bayesian} ecologist's shed},
	url = {https://onlinelibrary.wiley.com/doi/abs/10.1002/ece3.1346},
	doi = {10.1002/ece3.1346},
	abstract = {Despite benefits for precision, ecologists rarely use informative priors. One reason that ecologists may prefer vague priors is the perception that informative priors reduce accuracy. To date, no ecological study has empirically evaluated data-derived informative priors' effects on precision and accuracy. To determine the impacts of priors, we evaluated mortality models for tree species using data from a forest dynamics plot in Thailand. Half the models used vague priors, and the remaining half had informative priors. We found precision was greater when using informative priors, but effects on accuracy were more variable. In some cases, prior information improved accuracy, while in others, it was reduced. On average, models with informative priors were no more or less accurate than models without. Our analyses provide a detailed case study on the simultaneous effect of prior information on precision and accuracy and demonstrate that when priors are specified appropriately, they lead to greater precision without systematically reducing model accuracy.},
	language = {en},
	number = {1},
	urldate = {2024-03-29},
	journal = {Ecology and Evolution},
	author = {Morris, William K. and Vesk, Peter A. and McCarthy, Michael A. and Bunyavejchewin, Sarayudh and Baker, Patrick J.},
	year = {2015},
	keywords = {Ecological data, model precision, model validation, tree mortality},
	pages = {102--108},
}


@article{outhwaite_prior_2018,
	title = {Prior specification in {Bayesian} occupancy modelling improves analysis of species occurrence data},
	volume = {93},
	issn = {1470-160X},
	url = {https://www.sciencedirect.com/science/article/pii/S1470160X18303443},
	doi = {10.1016/j.ecolind.2018.05.010},
	abstract = {Multi-species biodiversity indicators are increasingly used to assess progress towards the 2020 ‘Aichi’ targets of the Convention on Biological Diversity. However, most multi-species indicators are biased towards a few well-studied taxa for which suitable abundance data are available. Consequently, many taxonomic groups are poorly represented in current measures of biodiversity change, particularly invertebrates. Alternative data sources, including opportunistic occurrence data, when analysed appropriately, can provide robust estimates of occurrence over time and increase the taxonomic coverage of such measures of population change. Occupancy modelling has been shown to produce robust estimates of species occurrence and trends through time. So far, this approach has concentrated on well-recorded taxa and performs poorly where recording intensity is low. Here, we show that the use of weakly informative priors in a Bayesian occupancy model framework greatly improves the precision of occurrence estimates associated with current model formulations when analysing low-intensity occurrence data, although estimated trends can be sensitive to the choice of prior when data are extremely sparse at either end of the recording period. Specifically, three variations of a Bayesian occupancy model, each with a different focus on information sharing among years, were compared using British ant data from the Bees, Wasps and Ants Recording Society and tested in a simulation experiment. Overall, the random walk model, which allows the sharing of information between the current and previous year, showed improved precision and low bias when estimating species occurrence and trends. The use of the model formulation described here will enable a greater range of datasets to be analysed, covering more taxa, which will significantly increase taxonomic representation of measures of biodiversity change.},
	urldate = {2024-03-29},
	journal = {Ecological Indicators},
	author = {Outhwaite, Charlotte L. and Chandler, Richard E. and Powney, Gary D. and Collen, Ben and Gregory, Richard D. and Isaac, Nick J. B.},
	month = oct,
	year = {2018},
	keywords = {Bayesian occupancy model, Biodiversity indicator, Biodiversity targets, Biological records, Multi-species indicator, Priors, Species occurrence},
	pages = {333--343},
}

@article{thomsen_monitoring_2012,
	title = {Monitoring endangered freshwater biodiversity using environmental {DNA}},
	volume = {21},
	copyright = {© 2011 Blackwell Publishing Ltd},
	issn = {1365-294X},
	url = {https://onlinelibrary.wiley.com/doi/abs/10.1111/j.1365-294X.2011.05418.x},
	doi = {10.1111/j.1365-294X.2011.05418.x},
	abstract = {Freshwater ecosystems are among the most endangered habitats on Earth, with thousands of animal species known to be threatened or already extinct. Reliable monitoring of threatened organisms is crucial for data-driven conservation actions but remains a challenge owing to nonstandardized methods that depend on practical and taxonomic expertise, which is rapidly declining. Here, we show that a diversity of rare and threatened freshwater animals—representing amphibians, fish, mammals, insects and crustaceans—can be detected and quantified based on DNA obtained directly from small water samples of lakes, ponds and streams. We successfully validate our findings in a controlled mesocosm experiment and show that DNA becomes undetectable within 2 weeks after removal of animals, indicating that DNA traces are near contemporary with presence of the species. We further demonstrate that entire faunas of amphibians and fish can be detected by high-throughput sequencing of DNA extracted from pond water. Our findings underpin the ubiquitous nature of DNA traces in the environment and establish environmental DNA as a tool for monitoring rare and threatened species across a wide range of taxonomic groups.},
	language = {en},
	number = {11},
	urldate = {2024-03-26},
	journal = {Molecular Ecology},
	author = {Thomsen, Philip Francis and Kielgast, Jos and Iversen, Lars L. and Wiuf, Carsten and Rasmussen, Morten and Gilbert, M. Thomas P and Orlando, Ludovic and Willerslev, Eske},
	year = {2012},
	keywords = {biological diversity, molecular detection, pyrosequencing, threatened species, wildlife conservation},
	pages = {2565--2573},
}


@article{campos-cerqueira_improving_2016,
	title = {Improving distribution data of threatened species by combining acoustic monitoring and occupancy modelling},
	volume = {7},
	copyright = {© 2016 The Authors. Methods in Ecology and Evolution © 2016 British Ecological Society},
	issn = {2041-210X},
	url = {https://onlinelibrary.wiley.com/doi/abs/10.1111/2041-210X.12599},
	doi = {10.1111/2041-210X.12599},
	abstract = {Conservation of threatened species relies on predictions about their spatial distribution; however, it is often difficult to detect species in the wild. The combination of acoustic monitoring to improve species detectability and statistical methods to account for false-negative detections can improve species distribution estimates. Here, we combine a novel automated species-specific identification approach with occupancy models that account for imperfect detectability to provide a more accurate species distribution map of the Elfin Woods Warbler Setophaga angelae, a rare, elusive and threatened bird species. We also compared three automated species identification/validation approaches to determine which approach provided occupancy estimates similar to manual validation of all recordings. Acoustic data were collected along three elevational gradients (95–1074 m a.s.l) in El Yunque National Forest, Puerto Rico. The detection matrices acquired through automated species-specific identification models and manual validations of all recordings were used to create occupancy models. Although this species has a wider distribution than previously reported, it depends on Palo Colorado forest cover and it mainly occurs between 600 and 900 m a.s.l. Unbiased and precise occupancy models were developed by using automated species identification models and only manually validating 4\% of the recordings. Our approach draws on the strength of two active areas of ecological research: acoustic monitoring and occupancy modelling. Our methods provide an effective and efficient way to translate the enormous amount of acoustic information collected with passive acoustic monitoring devices into meaningful ecological data that can be applied to understand and map the distribution of rare, elusive and threatened species.},
	language = {en},
	number = {11},
	urldate = {2024-03-26},
	journal = {Methods in Ecology and Evolution},
	author = {Campos-Cerqueira, Marconi and Aide, T. Mitchell},
	year = {2016},
	keywords = {automated species identification models, elusive, passive acoustic monitoring, portable recorders, rare, species distribution},
	pages = {1340--1348},
}


@article{sahu_environmental_2023,
	title = {Environmental {DNA} ({eDNA}): {Powerful} technique for biodiversity conservation},
	volume = {71},
	issn = {1617-1381},
	shorttitle = {Environmental {DNA} ({eDNA})},
	url = {https://www.sciencedirect.com/science/article/pii/S1617138122001984},
	doi = {10.1016/j.jnc.2022.126325},
	urldate = {2024-02-05},
	journal = {Journal for Nature Conservation},
	author = {Sahu, Ashish and Kumar, Neelesh and Pal Singh, Chandra and Singh, Mahender},
	month = feb,
	year = {2023},
	keywords = {Aquatic ecosystems, Biodiversity monitoring, Environmental DNA, Fish diversity, Metabarcoding},
	pages = {126325},
}

@article{ross_passive_2023,
	title = {Passive acoustic monitoring provides a fresh perspective on fundamental ecological questions},
	volume = {37},
	copyright = {© 2023 The Authors. Functional Ecology published by John Wiley \& Sons Ltd on behalf of British Ecological Society.},
	issn = {1365-2435},
	url = {https://onlinelibrary.wiley.com/doi/abs/10.1111/1365-2435.14275},
	doi = {10.1111/1365-2435.14275},
	language = {en},
	number = {4},
	urldate = {2024-02-05},
	journal = {Functional Ecology},
	author = {Ross, Samuel R. P.-J. and O'Connell, Darren P. and Deichmann, Jessica L. and Desjonquères, Camille and Gasc, Amandine and Phillips, Jennifer N. and Sethi, Sarab S. and Wood, Connor M. and Burivalova, Zuzana},
	year = {2023},
	keywords = {acoustic index, bioacoustics, biodiversity, ecoacoustics, sensor array, soundscape, spatiotemporal scale},
	pages = {959--975},
}


@article{welbourne_effectiveness_2015,
	title = {The effectiveness and cost of camera traps for surveying small reptiles and critical weight range mammals: {A} comparison with labour-intensive complementary methods},
	volume = {42},
	shorttitle = {The effectiveness and cost of camera traps for surveying small reptiles and critical weight range mammals},
	doi = {10.1071/WR15054},
	journal = {Wildlife Research},
	author = {Welbourne, Dustin and Macgregor, Christopher and Paull, David and Lindenmayer, David},
	month = sep,
	year = {2015},
	pages = {414--425},
}

@article{burton_review_2015,
	title = {{REVIEW}: {Wildlife} camera trapping: a review and recommendations for linking surveys to ecological processes},
	volume = {52},
	issn = {00218901},
	shorttitle = {{REVIEW}},
	url = {https://onlinelibrary.wiley.com/doi/10.1111/1365-2664.12432},
	doi = {10.1111/1365-2664.12432},
	language = {en},
	number = {3},
	urldate = {2023-05-25},
	journal = {Journal of Applied Ecology},
	author = {Burton, A. Cole and Neilson, Eric and Moreira, Dario and Ladle, Andrew and Steenweg, Robin and Fisher, Jason T. and Bayne, Erin and Boutin, Stan},
	month = jun,
	year = {2015},
	pages = {675--685},
}

@article{shonfield_autonomous_2017,
	title = {Autonomous recording units in avian ecological research: current use and future applications},
	volume = {12},
	issn = {1712-6568},
	shorttitle = {Autonomous recording units in avian ecological research},
	url = {http://www.ace-eco.org/vol12/iss1/art14/},
	doi = {10.5751/ACE-00974-120114},
	language = {en},
	number = {1},
	urldate = {2022-11-29},
	journal = {Avian Conservation and Ecology},
	author = {Shonfield, Julia and Bayne, Erin M.},
	year = {2017},
	pages = {14},
}

@article{hofmeester_framing_2019,
	title = {Framing pictures: {A} conceptual framework to identify and correct for biases in detection probability of camera traps enabling multi‐species comparison},
	volume = {9},
	issn = {2045-7758},
	shorttitle = {Framing pictures},
	url = {https://www.ncbi.nlm.nih.gov/pmc/articles/PMC6392353/},
	doi = {10.1002/ece3.4878},
	abstract = {Obtaining reliable species observations is of great importance in animal ecology and wildlife conservation. An increasing number of studies use camera traps (CTs) to study wildlife communities, and an increasing effort is made to make better use and reuse of the large amounts of data that are produced. It is in these circumstances that it becomes paramount to correct for the species‐ and study‐specific variation in imperfect detection within CTs. We reviewed the literature and used our own experience to compile a list of factors that affect CT detection of animals. We did this within a conceptual framework of six distinct scales separating out the influences of (a) animal characteristics, (b) CT specifications, (c) CT set‐up protocols, and (d) environmental variables. We identified 40 factors that can potentially influence the detection of animals by CTs at these six scales. Many of these factors were related to only a few overarching parameters. Most of the animal characteristics scale with body mass and diet type, and most environmental characteristics differ with season or latitude such that remote sensing products like NDVI could be used as a proxy index to capture this variation. Factors that influence detection at the microsite and camera scales are probably the most important in determining CT detection of animals. The type of study and specific research question will determine which factors should be corrected. Corrections can be done by directly adjusting the CT metric of interest or by using covariates in a statistical framework. Our conceptual framework can be used to design better CT studies and help when analyzing CT data. Furthermore, it provides an overview of which factors should be reported in CT studies to make them repeatable, comparable, and their data reusable. This should greatly improve the possibilities for global scale analyses of (reused) CT data.},
	number = {4},
	urldate = {2023-05-26},
	journal = {Ecology and Evolution},
	author = {Hofmeester, Tim R. and Cromsigt, Joris P. G. M. and Odden, John and Andrén, Henrik and Kindberg, Jonas and Linnell, John D. C.},
	month = jan,
	year = {2019},
	pages = {2320--2336},
}

@article{parsons_occupancy_2017,
	title = {Do occupancy or detection rates from camera traps reflect deer density?},
	volume = {98},
	issn = {0022-2372},
	url = {https://academic.oup.com/jmammal/article/98/6/1547/4430381},
	doi = {10.1093/jmammal/gyx128},
	abstract = {Abstract. Camera trapping is a powerful tool for studying mammal populations over large spatial scales. Density estimation using camera-trap data is a commonly},
	language = {en},
	number = {6},
	urldate = {2022-10-17},
	journal = {Journal of Mammalogy},
	author = {Parsons, Arielle Waldstein and Forrester, Tavis and McShea, William J. and Baker-Whatton, Megan C. and Millspaugh, Joshua J. and Kays, Roland},
	month = dec,
	year = {2017},
	pages = {1547--1557},
}


@article{nichols_multi-scale_2008,
	title = {Multi-scale occupancy estimation and modelling using multiple detection methods},
	volume = {45},
	issn = {1365-2664},
	url = {https://onlinelibrary.wiley.com/doi/abs/10.1111/j.1365-2664.2008.01509.x},
	doi = {10.1111/j.1365-2664.2008.01509.x},
	abstract = {1 Occupancy estimation and modelling based on detection–nondetection data provide an effective way of exploring change in a species’ distribution across time and space in cases where the species is not always detected with certainty. Today, many monitoring programmes target multiple species, or life stages within a species, requiring the use of multiple detection methods. When multiple methods or devices are used at the same sample sites, animals can be detected by more than one method. 2 We develop occupancy models for multiple detection methods that permit simultaneous use of data from all methods for inference about method-specific detection probabilities. Moreover, the approach permits estimation of occupancy at two spatial scales: the larger scale corresponds to species’ use of a sample unit, whereas the smaller scale corresponds to presence of the species at the local sample station or site. 3 We apply the models to data collected on two different vertebrate species: striped skunks Mephitis mephitis and red salamanders Pseudotriton ruber. For striped skunks, large-scale occupancy estimates were consistent between two sampling seasons. Small-scale occupancy probabilities were slightly lower in the late winter/spring when skunks tend to conserve energy, and movements are limited to males in search of females for breeding. There was strong evidence of method-specific detection probabilities for skunks. As anticipated, large- and small-scale occupancy areas completely overlapped for red salamanders. The analyses provided weak evidence of method-specific detection probabilities for this species. 4 Synthesis and applications. Increasingly, many studies are utilizing multiple detection methods at sampling locations. The modelling approach presented here makes efficient use of detections from multiple methods to estimate occupancy probabilities at two spatial scales and to compare detection probabilities associated with different detection methods. The models can be viewed as another variation of Pollock's robust design and may be applicable to a wide variety of scenarios where species occur in an area but are not always near the sampled locations. The estimation approach is likely to be especially useful in multispecies conservation programmes by providing efficient estimates using multiple detection devices and by providing device-specific detection probability estimates for use in survey design.},
	language = {en},
	number = {5},
	urldate = {2024-01-31},
	journal = {Journal of Applied Ecology},
	author = {Nichols, James D. and Bailey, Larissa L. and O’Connell Jr., Allan F. and Talancy, Neil W. and Campbell Grant, Evan H. and Gilbert, Andrew T. and Annand, Elizabeth M. and Husband, Thomas P. and Hines, James E.},
	year = {2008},
	keywords = {Pollock's robust design, detection probability, model, occupancy, red salamander, sampling, striped skunk},
	pages = {1321--1329},
}


@article{thompson_use_2007,
	title = {Use of {Site} {Occupancy} {Models} to {Estimate} {Prevalence} of {Myxobolus} cerebralis {Infection} in {Trout}},
	volume = {19},
	copyright = {© 2007 American Fisheries Society},
	issn = {1548-8667},
	url = {https://onlinelibrary.wiley.com/doi/abs/10.1577/H06-016.1},
	doi = {10.1577/H06-016.1},
	language = {en},
	number = {1},
	urldate = {2024-01-31},
	journal = {Journal of Aquatic Animal Health},
	author = {Thompson, Kevin G.},
	year = {2007},
	pages = {8--13},
}

@article{lachish_site-occupancy_2012,
	title = {Site-occupancy modelling as a novel framework for assessing test sensitivity and estimating wildlife disease prevalence from imperfect diagnostic tests},
	volume = {3},
	copyright = {© 2011 The Authors. Methods in Ecology and Evolution © 2011 British Ecological Society},
	issn = {2041-210X},
	url = {https://onlinelibrary.wiley.com/doi/abs/10.1111/j.2041-210X.2011.00156.x},
	doi = {10.1111/j.2041-210X.2011.00156.x},
	language = {en},
	number = {2},
	urldate = {2024-01-26},
	journal = {Methods in Ecology and Evolution},
	author = {Lachish, Shelly and Gopalaswamy, Arjun M. and Knowles, Sarah C. L. and Sheldon, Ben C.},
	year = {2012},
	keywords = {Cyanistes caeruleus, Plasmodium, imperfect diagnostic tests, pathogen detection, pathogen load, population prevalence, qPCR, site-occupancy modelling, test sensitivity, wildlife disease},
	pages = {339--348},
}

@article{tyre_improving_2003,
	title = {Improving {Precision} and {Reducing} {Bias} in {Biological} {Surveys}: {Estimating} {False}-{Negative} {Error} {Rates}},
	volume = {13},
	copyright = {© 2003 by the Ecological Society of America},
	issn = {1939-5582},
	shorttitle = {Improving {Precision} and {Reducing} {Bias} in {Biological} {Surveys}},
	url = {https://onlinelibrary.wiley.com/doi/abs/10.1890/02-5078},
	doi = {10.1890/02-5078},
	language = {en},
	number = {6},
	urldate = {2024-01-24},
	journal = {Ecological Applications},
	author = {Tyre, Andrew J. and Tenhumberg, Brigitte and Field, Scott A. and Niejalke, Darren and Parris, Kirsten and Possingham, Hugh P.},
	year = {2003},
	keywords = {biological surveys, false-negative errors, habitat effects, presence–absence data, zero-inflated binomial (ZIB) model},
	pages = {1790--1801},
}

@article{hunter_environmental_2015,
	title = {Environmental {DNA} ({eDNA}) {Sampling} {Improves} {Occurrence} and {Detection} {Estimates} of {Invasive} {Burmese} {Pythons}},
	volume = {10},
	issn = {1932-6203},
	url = {https://journals.plos.org/plosone/article?id=10.1371/journal.pone.0121655},
	doi = {10.1371/journal.pone.0121655},
	abstract = {Environmental DNA (eDNA) methods are used to detect DNA that is shed into the aquatic environment by cryptic or low density species. Applied in eDNA studies, occupancy models can be used to estimate occurrence and detection probabilities and thereby account for imperfect detection. However, occupancy terminology has been applied inconsistently in eDNA studies, and many have calculated occurrence probabilities while not considering the effects of imperfect detection. Low detection of invasive giant constrictors using visual surveys and traps has hampered the estimation of occupancy and detection estimates needed for population management in southern Florida, USA. Giant constrictor snakes pose a threat to native species and the ecological restoration of the Florida Everglades. To assist with detection, we developed species-specific eDNA assays using quantitative PCR (qPCR) for the Burmese python (Python molurus bivittatus), Northern African python (P. sebae), boa constrictor (Boa constrictor), and the green (Eunectes murinus) and yellow anaconda (E. notaeus). Burmese pythons, Northern African pythons, and boa constrictors are established and reproducing, while the green and yellow anaconda have the potential to become established. We validated the python and boa constrictor assays using laboratory trials and tested all species in 21 field locations distributed in eight southern Florida regions. Burmese python eDNA was detected in 37 of 63 field sampling events; however, the other species were not detected. Although eDNA was heterogeneously distributed in the environment, occupancy models were able to provide the first estimates of detection probabilities, which were greater than 91\%. Burmese python eDNA was detected along the leading northern edge of the known population boundary. The development of informative detection tools and eDNA occupancy models can improve conservation efforts in southern Florida and support more extensive studies of invasive constrictors. Generic sampling design and terminology are proposed to standardize and clarify interpretations of eDNA-based occupancy models.},
	language = {en},
	number = {4},
	urldate = {2023-11-06},
	journal = {PLOS ONE},
	author = {Hunter, Margaret E. and Oyler-McCance, Sara J. and Dorazio, Robert M. and Fike, Jennifer A. and Smith, Brian J. and Hunter, Charles T. and Reed, Robert N. and Hart, Kristen M.},
	year = {2015},
	keywords = {Birds, DNA extraction, DNA replication, Florida, Invasive species, Polymerase chain reaction, Pythons, Snakes},
	pages = {e0121655},
}


@article{willoughby_importance_2016,
	title = {The importance of including imperfect detection models in {eDNA} experimental design},
	volume = {16},
	copyright = {© 2016 John Wiley \& Sons Ltd},
	issn = {1755-0998},
	url = {https://onlinelibrary.wiley.com/doi/abs/10.1111/1755-0998.12531},
	language = {en},
	number = {4},
	urldate = {2023-11-06},
	journal = {Molecular Ecology Resources},
	author = {Willoughby, Janna R. and Wijayawardena, Bhagya K. and Sundaram, Mekala and Swihart, Robert K. and DeWoody, J. Andrew},
	year = {2016},
	keywords = {environmental DNA, occupancy model, probability of detection, sampling effort},
	pages = {837--844},
}

@article{schmidt_site_2013,
	title = {Site occupancy models in the analysis of environmental {DNA} presence/absence surveys: a case study of an emerging amphibian pathogen},
	volume = {4},
	copyright = {© 2013 The Authors. Methods in Ecology and Evolution © 2013 British Ecological Society},
	issn = {2041-210X},
	shorttitle = {Site occupancy models in the analysis of environmental {DNA} presence/absence surveys},
	url = {https://onlinelibrary.wiley.com/doi/abs/10.1111/2041-210X.12052},
	doi = {10.1111/2041-210X.12052},
	language = {en},
	number = {7},
	urldate = {2023-11-06},
	journal = {Methods in Ecology and Evolution},
	author = {Schmidt, Benedikt R. and Kéry, Marc and Ursenbacher, Sylvain and Hyman, Oliver J. and Collins, James P.},
	year = {2013},
	keywords = {detection probability, environmental DNA, monitoring, site occupancy model, survey},
	pages = {646--653},
}

@article{doi_evaluation_2019,
	title = {Evaluation of detection probabilities at the water-filtering and initial {PCR} steps in environmental {DNA} metabarcoding using a multispecies site occupancy model},
	volume = {9},
	copyright = {2019 The Author(s)},
	issn = {2045-2322},
	url = {https://www.nature.com/articles/s41598-019-40233-1},
	doi = {10.1038/s41598-019-40233-1},
	language = {en},
	number = {1},
	urldate = {2023-11-06},
	journal = {Scientific Reports},
	author = {Doi, Hideyuki and Fukaya, Keiichi and Oka, Shin-ichiro and Sato, Keiichi and Kondoh, Michio and Miya, Masaki},
	month = mar,
	year = {2019},
	keywords = {Community ecology, Molecular ecology},
	pages = {3581},
}

@article{gold_comparison_2023,
	title = {A comparison of biomonitoring methodologies for surf zone fish communities},
	volume = {18},
	issn = {1932-6203},
	url = {https://journals.plos.org/plosone/article?id=10.1371/journal.pone.0260903},
	doi = {10.1371/journal.pone.0260903},
	language = {en},
	number = {6},
	urldate = {2023-10-16},
	journal = {PLOS ONE},
	author = {Gold, Zachary and Koch, McKenzie Q. and Schooler, Nicholas K. and Emery, Kyle A. and Dugan, Jenifer E. and Miller, Robert J. and Page, Henry M. and Schroeder, Donna M. and Hubbard, David M. and Madden, Jessica R. and Whitaker, Stephen G. and Barber, Paul H.},
	year = {2023},
	keywords = {Beaches, Biodiversity, Coastal ecosystems, Ecosystems, Marine ecosystems, Marine fish, Survey methods, Taxonomy},
	pages = {e0260903},
}

@article{cole_determining_2010,
	title = {Determining the parametric structure of models},
	volume = {228},
	issn = {0025-5564},
	url = {https://www.sciencedirect.com/science/article/pii/S0025556410001240},
	doi = {10.1016/j.mbs.2010.08.004},
	number = {1},
	urldate = {2023-10-13},
	journal = {Mathematical Biosciences},
	author = {Cole, D. J. and Morgan, B. J. T. and Titterington, D. M.},
	month = nov,
	year = {2010},
	keywords = {Derivative matrix, Exhaustive summaries, Global identifiability, Jacobian matrix, Local identifiability, Parameter redundancy},
	pages = {16--30},
}

@article{mcclintock_seeking_2010,
	title = {Seeking a second opinion: uncertainty in disease ecology: {Uncertainty} in disease ecology},
	volume = {13},
	issn = {1461023X, 14610248},
	shorttitle = {Seeking a second opinion},
	url = {https://onlinelibrary.wiley.com/doi/10.1111/j.1461-0248.2010.01472.x},
	doi = {10.1111/j.1461-0248.2010.01472.x},
	language = {en},
	number = {6},
	urldate = {2023-10-13},
	journal = {Ecology Letters},
	author = {McClintock, Brett T. and Nichols, James D. and Bailey, Larissa L. and MacKenzie, Darryl I. and Kendall, William. L. and Franklin, Alan B.},
	month = apr,
	year = {2010},
	pages = {659--674},
}


@article{mcclintock_unmodeled_2010,
	title = {Unmodeled observation error induces bias when inferring patterns and dynamics of species occurrence via aural detections},
	volume = {91},
	copyright = {© 2010 by the Ecological Society of America},
	issn = {1939-9170},
	url = {https://onlinelibrary.wiley.com/doi/abs/10.1890/09-1287.1},
	doi = {10.1890/09-1287.1},
	language = {en},
	number = {8},
	urldate = {2023-10-13},
	journal = {Ecology},
	author = {McClintock, Brett T. and Bailey, Larissa L. and Pollock, Kenneth H. and Simons, Theodore R.},
	year = {2010},
	keywords = {auditory detection, colonization, detection probability, false negative, false positive, imperfect detection, local extinction, measurement error, monitoring, observation error, site occupancy, species occurrence},
	pages = {2446--2454},
}

@article{da_silva_neto_integrating_2020,
	title = {Integrating species distribution and occupancy modeling to study hellbender ({Cryptobranchus} alleganiensis) occurrence based on {eDNA} surveys},
	volume = {251},
	issn = {00063207},
	url = {https://linkinghub.elsevier.com/retrieve/pii/S0006320720308454},
	doi = {10.1016/j.biocon.2020.108787},
	language = {en},
	urldate = {2023-10-13},
	journal = {Biological Conservation},
	author = {Da Silva Neto, Jeronimo G. and Sutton, William B. and Spear, Stephen F. and Freake, Michael J. and Kéry, Marc and Schmidt, Benedikt R.},
	month = nov,
	year = {2020},
	pages = {108787},
}


@article{gimenez_methods_2004,
    author = {Gimenez, O. and Viallefont, A. and Catchpole, E. A. and Choquet, R. and Morgan, B. J. T.},
    title = {Methods for investigating parameter redundancy},
    year = {2004},
    volume = {27.1},
    pages = {561--572},
    journal = {Animal Biodiversity and Conservation},
    keywords = {Mark-recapture data, Mark-recovery data, Profile-likelihood, Analytical-numerical method, Symbolic algebra software}
}

@article{griffin_modelling_2020,
	title = {Modelling environmental {DNA} data; {Bayesian} variable selection accounting for false positive and false negative errors},
	volume = {69},
	copyright = {© 2019 Royal Statistical Society},
	issn = {1467-9876},
	url = {https://onlinelibrary.wiley.com/doi/abs/10.1111/rssc.12390},
	doi = {10.1111/rssc.12390},
	language = {en},
	number = {2},
	urldate = {2023-09-18},
	journal = {Journal of the Royal Statistical Society: Series C (Applied Statistics)},
	author = {Griffin, Jim E. and Matechou, Eleni and Buxton, Andrew S. and Bormpoudakis, Dimitrios and Griffiths, Richard A.},
	year = {2020},
	keywords = {Informative prior distributions, Known presences, Likelihood symmetries, Logistic regression, Occupancy probability, Pólya–gamma scheme},
	pages = {377--392},
}

@article{belmont_new_2022,
	title = {A new statistical approach for identifying rare species under imperfect detection},
	volume = {28},
	copyright = {© 2022 The Authors. Diversity and Distributions published by John Wiley \& Sons Ltd.},
	issn = {1472-4642},
	url = {https://onlinelibrary.wiley.com/doi/abs/10.1111/ddi.13495},
	doi = {10.1111/ddi.13495},
	language = {en},
	number = {5},
	urldate = {2023-09-13},
	journal = {Diversity and Distributions},
	author = {Belmont, Jafet and Miller, Claire and Scott, Marian and Wilkie, Craig},
	year = {2022},
	keywords = {Odonata, classification, community ecology, detectability, occupancy model, rare species},
	pages = {882--893},
}

@article{mckibben_model_2023,
	title = {To model or not to model: false positive detection error in camera surveys},
	volume = {87},
	copyright = {© 2023 The Wildlife Society},
	issn = {1937-2817},
	shorttitle = {To model or not to model},
	url = {https://onlinelibrary.wiley.com/doi/abs/10.1002/jwmg.22365},
	doi = {10.1002/jwmg.22365},
	language = {en},
	number = {3},
	urldate = {2023-09-12},
	journal = {The Journal of Wildlife Management},
	author = {McKibben, Fiona E. and Abadi, Fitsum and Frey, Jennifer K.},
	year = {2023},
	keywords = {Neotamias minimus atristriatus, Peñasco least chipmunk, detection error, false positive error, gray-footed chipmunk, imperfect detection, misidentification, occupancy, remote camera, survey},
	pages = {e22365},
}

@article{ferguson_occupancy_2015,
	title = {Occupancy models for data with false positive and false negative errors and heterogeneity across sites and surveys},
	volume = {6},
	copyright = {© 2015 The Authors. Methods in Ecology and Evolution © 2015 British Ecological Society},
	issn = {2041-210X},
	url = {https://onlinelibrary.wiley.com/doi/abs/10.1111/2041-210X.12442},
	doi = {10.1111/2041-210X.12442},
	language = {en},
	number = {12},
	urldate = {2023-09-11},
	journal = {Methods in Ecology and Evolution},
	author = {Ferguson, Paige F.B. and Conroy, Michael J. and Hepinstall-Cymerman, Jeffrey},
	year = {2015},
	keywords = {Monte Carlo simulation, confirmed detection, detection probability, hierarchical Bayesian model, imperfect detection, informative prior, observation, occupancy model, phantom species, scenario},
	pages = {1395--1406},
}


@article{lonsinger_efficacy_2023,
	title = {Efficacy of machine learning image classification for automated occupancy‐based monitoring},
	issn = {2056-3485, 2056-3485},
	url = {https://zslpublications.onlinelibrary.wiley.com/doi/10.1002/rse2.356},
	doi = {10.1002/rse2.356},
	language = {en},
	urldate = {2023-08-22},
	journal = {Remote Sensing in Ecology and Conservation},
	author = {Lonsinger, Robert C. and Dart, Marlin M. and Larsen, Randy T. and Knight, Robert N.},
	month = jul,
	year = {2023},
	pages = {56-71},
}


@article{gimenez_tradeoff_2022,
  author = {Olivier Gimenez and Maëlis Kervellec and Jean-Baptiste Fanjul and Anna Chaine and Lucile Marescot and Yoann Bollet and Christophe Duchamp},
  publisher = {Société Française de Statistique},
  title = {Trade-Off Between Deep Learning for Species Identification and Inference about Predator-Prey Co-Occurrence},
  journal = {Computo},
  year = {2022},
  url = {https://computo.sfds.asso.fr/published-202204-deeplearning-occupancy-lynx},
  doi = {10.57750/yfm2-5f45},
  issn = {2824-7795},
  langid = {en},
}


@article{guillera-arroita_dealing_2017,
	title = {Dealing with false-positive and false-negative errors about species occurrence at multiple levels},
	volume = {8},
	copyright = {© 2017 The Authors. Methods in Ecology and Evolution © 2017 British Ecological Society},
	issn = {2041-210X},
	url = {https://onlinelibrary.wiley.com/doi/abs/10.1111/2041-210X.12743},
	doi = {10.1111/2041-210X.12743},
	language = {en},
	number = {9},
	urldate = {2023-06-02},
	journal = {Methods in Ecology and Evolution},
	author = {Guillera-Arroita, Gurutzeta and Lahoz-Monfort, José Joaquín and van Rooyen, Anthony R. and Weeks, Andrew R. and Tingley, Reid},
	year = {2017},
	keywords = {detectability, environmental DNA, identifiability, imperfect detection, monitoring, multiple stages, sensitivity, species occupancy, specificity},
	pages = {1081--1091},
}


@article{swanson_snapshot_2015,
	title = {Snapshot {Serengeti}, high-frequency annotated camera trap images of 40 mammalian species in an {African} savanna},
	volume = {2},
	copyright = {2015 The Author(s)},
	issn = {2052-4463},
	url = {https://www.nature.com/articles/sdata201526},
	doi = {10.1038/sdata.2015.26},
	language = {en},
	number = {1},
	urldate = {2023-03-31},
	journal = {Scientific Data},
	author = {Swanson, Alexandra and Kosmala, Margaret and Lintott, Chris and Simpson, Robert and Smith, Arfon and Packer, Craig},
	month = jun,
	year = {2015},
	keywords = {Biodiversity, Community ecology, Population dynamics},
	pages = {150026},
}


@article{velez_evaluation_2023,
	title = {An evaluation of platforms for processing camera-trap data using artificial intelligence},
	volume = {14},
	issn = {2041-210X},
	url = {https://onlinelibrary.wiley.com/doi/abs/10.1111/2041-210X.14044},
	doi = {10.1111/2041-210X.14044},
	language = {en},
	number = {2},
	urldate = {2023-03-15},
	journal = {Methods in Ecology and Evolution},
	author = {Vélez, Juliana and McShea, William and Shamon, Hila and Castiblanco-Camacho, Paula J. and Tabak, Michael A. and Chalmers, Carl and Fergus, Paul and Fieberg, John},
	year = {2023},
	keywords = {artificial intelligence, camera traps, computer vision, data processing, deep learning, image classification, remote sensing, review},
	pages = {459--477},
}

@techreport{augustine_towards_2023,
	type = {preprint},
	title = {Towards estimating marine wildlife abundance using aerial surveys and deep learning with hierarchical classifications subject to error},
	url = {http://biorxiv.org/lookup/doi/10.1101/2023.02.20.529272},
	language = {en},
	urldate = {2023-02-23},
	institution = {Ecology},
	author = {Augustine, Ben C and Koneff, Mark D and Pickens, Bradley A. and Royle, J. Andrew},
    Publisher = {bioRxiv},
	month = feb,
	year = {2023},
	doi = {10.1101/2023.02.20.529272},
}


@article{wrege_acoustic_2017,
	title = {Acoustic monitoring for conservation in tropical forests: examples from forest elephants},
	volume = {8},
	issn = {2041-210X},
	shorttitle = {Acoustic monitoring for conservation in tropical forests},
	url = {https://onlinelibrary.wiley.com/doi/abs/10.1111/2041-210X.12730},
	doi = {10.1111/2041-210X.12730},
	language = {en},
	number = {10},
	urldate = {2023-01-17},
	journal = {Methods in Ecology and Evolution},
	author = {Wrege, Peter H. and Rowland, Elizabeth D. and Keen, Sara and Shiu, Yu},
	year = {2017},
	keywords = {Loxodonta cyclotis, anthropogenic impacts, anti-poaching, bioacoustics, cryptic species, density estimate, law enforcement monitoring, passive acoustic monitoring, signal detection, smart},
	pages = {1292--1301},
}


@article{duggan_approach_2021,
	title = {An approach to rapid processing of camera trap images with minimal human input},
	volume = {11},
	issn = {2045-7758},
	url = {https://onlinelibrary.wiley.com/doi/abs/10.1002/ece3.7970},
	doi = {10.1002/ece3.7970},
	language = {en},
	number = {17},
	urldate = {2023-01-06},
	journal = {Ecology and Evolution},
	author = {Duggan, Matthew T. and Groleau, Melissa F. and Shealy, Ethan P. and Self, Lillian S. and Utter, Taylor E. and Waller, Matthew M. and Hall, Bryan C. and Stone, Chris G. and Anderson, Layne L. and Mousseau, Timothy A.},
	year = {2021},
	keywords = {camera trap, deep learning, neural network, transfer learning, wildlife ecology},
	pages = {12051--12063},
}


@article{findlay_component_2020,
	title = {Component processes of detection probability in camera-trap studies: understanding the occurrence of false-negatives},
	volume = {65},
	issn = {2199-241X},
	shorttitle = {Component processes of detection probability in camera-trap studies},
	url = {https://doi.org/10.1007/s13364-020-00478-y},
	doi = {10.1007/s13364-020-00478-y},
	language = {en},
	number = {2},
	urldate = {2022-12-21},
	journal = {Mammal Research},
	author = {Findlay, Melanie A. and Briers, Robert A. and White, Patrick J. C.},
	month = apr,
	year = {2020},
	keywords = {Camera-trap, Data quality, Detection, False-negatives},
	pages = {167--180},
}


@article{chambert_modeling_2015,
	title = {Modeling false positive detections in species occurrence data under different study designs},
	volume = {96},
	issn = {1939-9170},
	url = {https://onlinelibrary.wiley.com/doi/abs/10.1890/14-1507.1},
	doi = {10.1890/14-1507.1},
	language = {en},
	number = {2},
	urldate = {2022-12-06},
	journal = {Ecology},
	author = {Chambert, Thierry and Miller, David A. W. and Nichols, James D.},
	year = {2015},
	keywords = {detection error, false negative, false positive, misidentification, observation error, occupancy modeling, species distribution modeling},
	pages = {332--339},
}


@article{barre_accounting_2019,
	title = {Accounting for automated identification errors in acoustic surveys},
	volume = {10},
	issn = {2041-210X},
	url = {https://onlinelibrary.wiley.com/doi/abs/10.1111/2041-210X.13198},
	doi = {10.1111/2041-210X.13198},
	language = {en},
	number = {8},
	urldate = {2022-11-29},
	journal = {Methods in Ecology and Evolution},
	author = {Barré, Kévin and Le Viol, Isabelle and Julliard, Romain and Pauwels, Julie and Newson, Stuart E. and Julien, Jean-François and Claireau, Fabien and Kerbiriou, Christian and Bas, Yves},
	year = {2019},
	keywords = {bioacoustic, cautious threshold, chiroptera, error rate, false positives, passive acoustic monitoring, semi-automated identification, survey methods},
	pages = {1171--1188},
}

@article{spiers_estimating_2022,
	title = {Estimating species misclassification with occupancy dynamics and encounter rates: {A} semi‐supervised, individual‐level approach},
	volume = {13},
	issn = {2041-210X, 2041-210X},
	shorttitle = {Estimating species misclassification with occupancy dynamics and encounter rates},
	url = {https://onlinelibrary.wiley.com/doi/10.1111/2041-210X.13858},
	doi = {10.1111/2041-210X.13858},
	language = {en},
	number = {7},
	urldate = {2022-11-04},
	journal = {Methods in Ecology and Evolution},
	author = {Spiers, Anna I. and Royle, J. Andrew and Torrens, Christa L. and Joseph, Maxwell B.},
	month = jul,
	year = {2022},
	pages = {1528--1539},
}


@article{miller_improving_2011,
	title = {Improving occupancy estimation when two types of observational error occur: non-detection and species misidentification},
	volume = {92},
	issn = {1939-9170},
	shorttitle = {Improving occupancy estimation when two types of observational error occur},
	url = {https://onlinelibrary.wiley.com/doi/abs/10.1890/10-1396.1},
	doi = {10.1890/10-1396.1},
	language = {en},
	number = {7},
	urldate = {2022-11-04},
	journal = {Ecology},
	author = {Miller, David A. and Nichols, James D. and McClintock, Brett T. and Grant, Evan H. Campbell and Bailey, Larissa L. and Weir, Linda A.},
	year = {2011},
	keywords = {Lithobates spp., anuran censuses, call surveys, false positive detection, misclassification, misidentification, multiple states, presence–absence, proportion area occupied, site occupancy, species occurrence},
	pages = {1422--1428},
}

@article{mackenzie_estimating_2002,
	title = {Estimating site occupancy rates when detection probabilities are less than one},
	volume = {83},
	issn = {0012-9658},
	url = {http://doi.wiley.com/10.1890/0012-9658(2002)083[2248:ESORWD]2.0.CO;2},
	doi = {10.1890/0012-9658(2002)083[2248:ESORWD]2.0.CO;2},
	language = {en},
	number = {8},
	urldate = {2022-10-13},
	journal = {Ecology},
	author = {MacKenzie, Darryl I. and Nichols, James D. and Lachman, Gideon B. and Droege, Sam and Andrew Royle, J. and Langtimm, Catherine A.},
	month = aug,
	year = {2002},
	pages = {2248--2255},
}

@article{tabak_machine_2019,
	title = {Machine learning to classify animal species in camera trap images: {Applications} in ecology},
	volume = {10},
	issn = {2041-210X},
	shorttitle = {Machine learning to classify animal species in camera trap images},
	url = {https://onlinelibrary.wiley.com/doi/abs/10.1111/2041-210X.13120},
	doi = {10.1111/2041-210X.13120},
	language = {en},
	number = {4},
	urldate = {2022-10-05},
	journal = {Methods in Ecology and Evolution},
	author = {Tabak, Michael A. and Norouzzadeh, Mohammad S. and Wolfson, David W. and Sweeney, Steven J. and Vercauteren, Kurt C. and Snow, Nathan P. and Halseth, Joseph M. and Di Salvo, Paul A. and Lewis, Jesse S. and White, Michael D. and Teton, Ben and Beasley, James C. and Schlichting, Peter E. and Boughton, Raoul K. and Wight, Bethany and Newkirk, Eric S. and Ivan, Jacob S. and Odell, Eric A. and Brook, Ryan K. and Lukacs, Paul M. and Moeller, Anna K. and Mandeville, Elizabeth G. and Clune, Jeff and Miller, Ryan S.},
	year = {2019},
	keywords = {artificial intelligence, camera trap, convolutional neural network, deep neural networks, image classification, machine learning, r package, remote sensing},
	pages = {585--590},
}

@article{tabak_improving_2020,
	title = {Improving the accessibility and transferability of machine learning algorithms for identification of animals in camera trap images: {MLWIC2}},
	volume = {10},
	issn = {2045-7758},
	shorttitle = {Improving the accessibility and transferability of machine learning algorithms for identification of animals in camera trap images},
	url = {https://onlinelibrary.wiley.com/doi/abs/10.1002/ece3.6692},
	doi = {10.1002/ece3.6692},
	language = {en},
	number = {19},
	urldate = {2022-10-05},
	journal = {Ecology and Evolution},
	author = {Tabak, Michael A. and Norouzzadeh, Mohammad S. and Wolfson, David W. and Newton, Erica J. and Boughton, Raoul K. and Ivan, Jacob S. and Odell, Eric A. and Newkirk, Eric S. and Conrey, Reesa Y. and Stenglein, Jennifer and Iannarilli, Fabiola and Erb, John and Brook, Ryan K. and Davis, Amy J. and Lewis, Jesse and Walsh, Daniel P. and Beasley, James C. and VerCauteren, Kurt C. and Clune, Jeff and Miller, Ryan S.},
	year = {2020},
	keywords = {R package, computer vision, deep convolutional neural networks, image classification, machine learning, motion-activated camera, remote sensing, species identification},
	pages = {10374--10383},
}


@article{norouzzadeh_deep_2021,
	title = {A deep active learning system for species identification and counting in camera trap images},
	volume = {12},
	issn = {2041-210X},
	url = {https://onlinelibrary.wiley.com/doi/abs/10.1111/2041-210X.13504},
	doi = {10.1111/2041-210X.13504},
	language = {en},
	number = {1},
	urldate = {2022-10-05},
	journal = {Methods in Ecology and Evolution},
	author = {Norouzzadeh, Mohammad Sadegh and Morris, Dan and Beery, Sara and Joshi, Neel and Jojic, Nebojsa and Clune, Jeff},
	year = {2021},
	keywords = {active learning, camera trap images, computer vision, deep learning, deep neural networks},
	pages = {150--161},
}

@article{royle_generalized_2006,
	title = {Generalized {Site} {Occupancy} {Models} {Allowing} for {False} {Positive} and {False} {Negative} {Errors}},
	volume = {87},
	issn = {1939-9170},
	url = {http://onlinelibrary.wiley.com/doi/abs/10.1890/0012-9658%282006%2987%5B835%3AGSOMAF%5D2.0.CO%3B2},
	doi = {10.1890/0012-9658(2006)87[835:GSOMAF]2.0.CO;2},
	language = {en},
	number = {4},
	urldate = {2022-11-04},
	journal = {Ecology},
	author = {Royle, J. Andrew and Link, William A.},
	year = {2006},
	keywords = {finite mixture, heterogeneous detection probability, latent class model, multinomial misclassification, multinomial mixtures, nondetection bias, occurrence probability, proportion of area occupied},
	pages = {835--841},
}
