## Supplementary figures and images for "Using informative priors to account for identifiability issues in occupancy models with identification errors"

### badge_PCI_Ecology.png

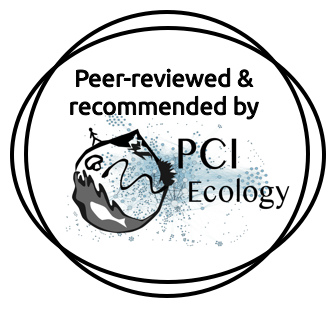

### figure1.jpeg

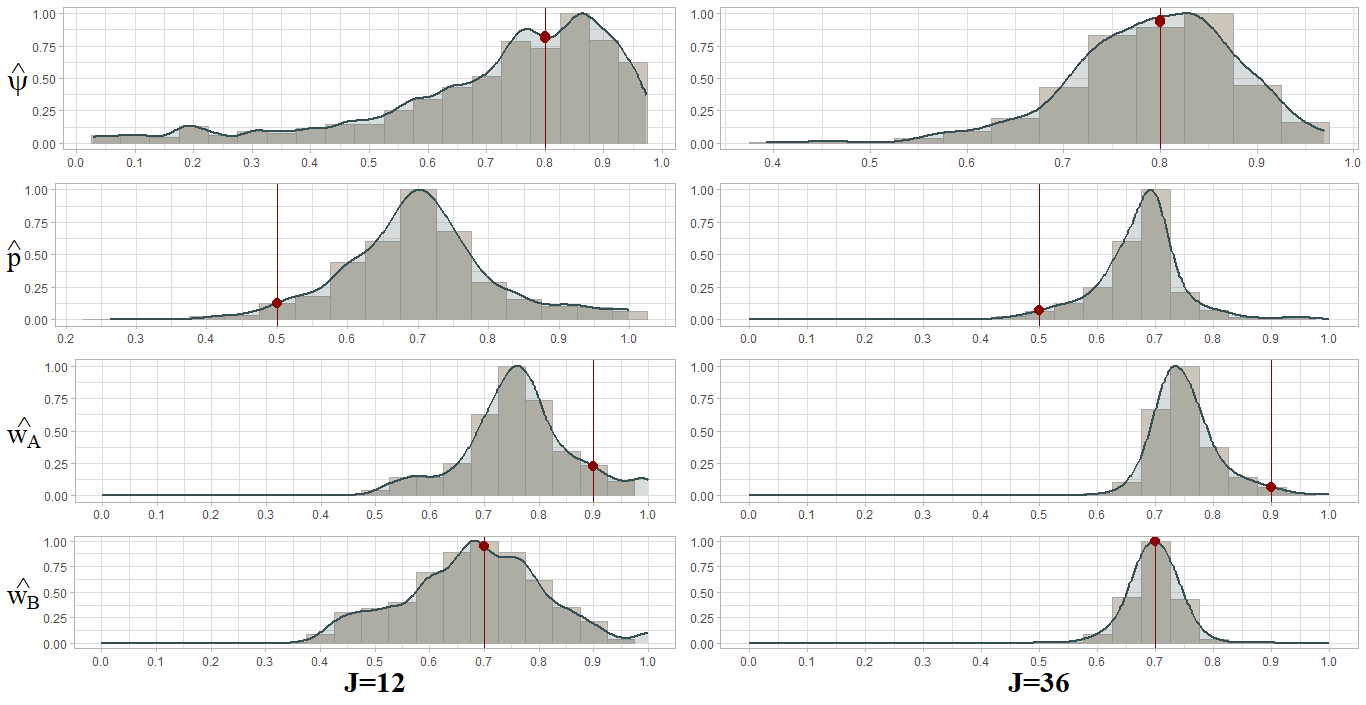

### figure2.jpeg

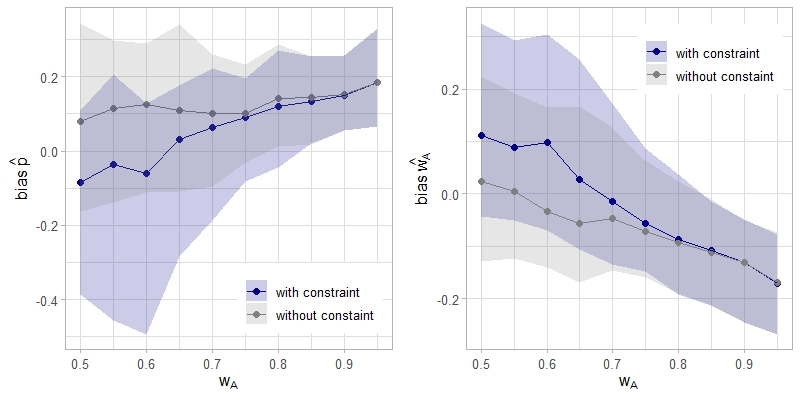

### figure3.png

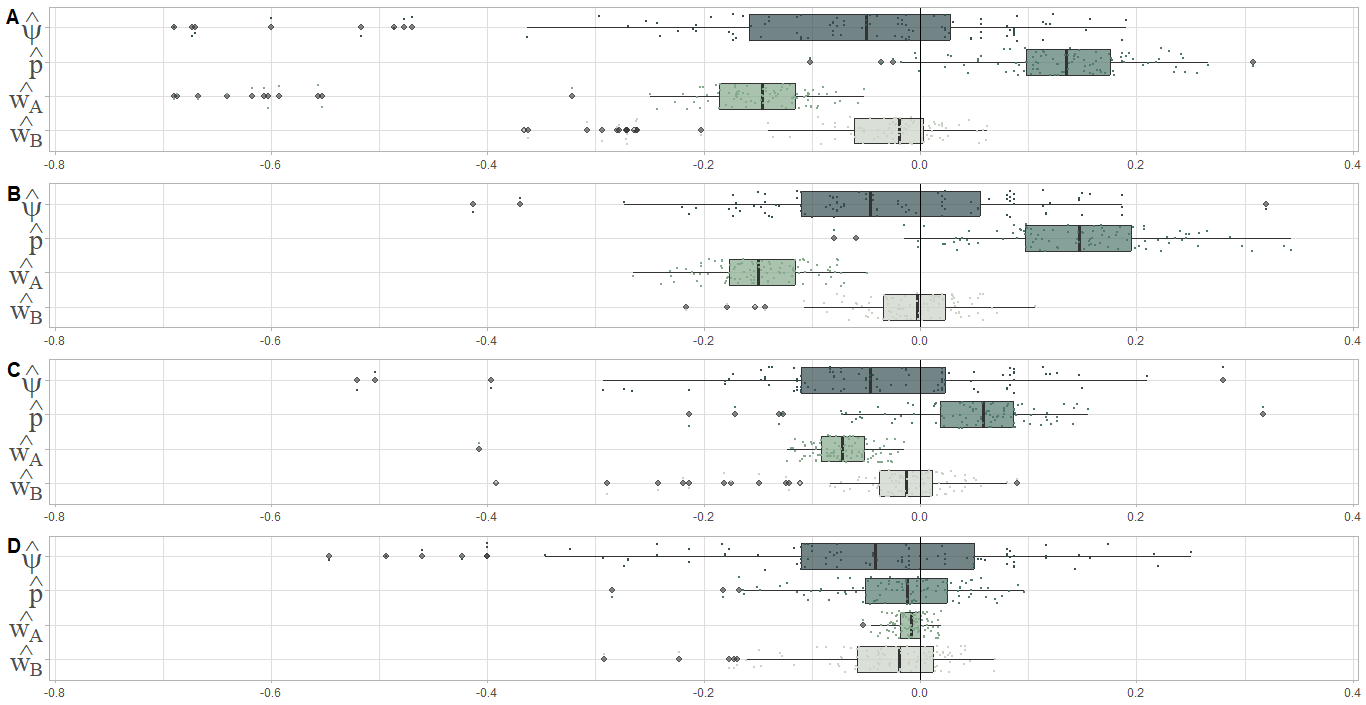

### figureA1.png

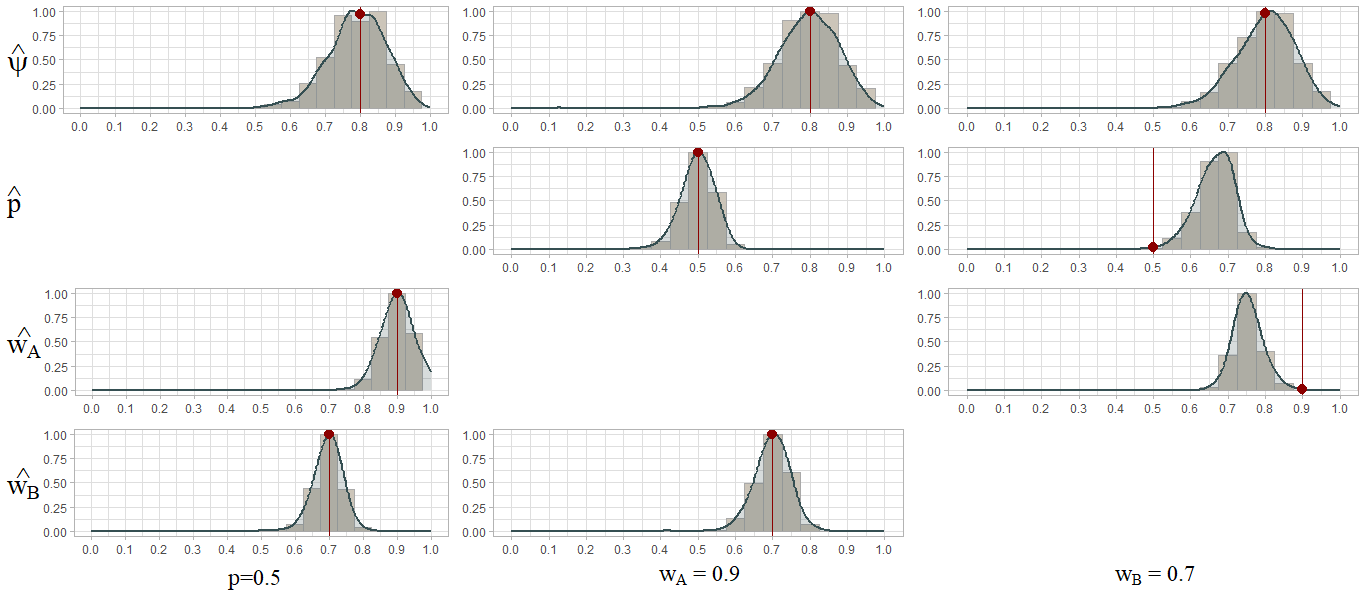

### figureA2.png

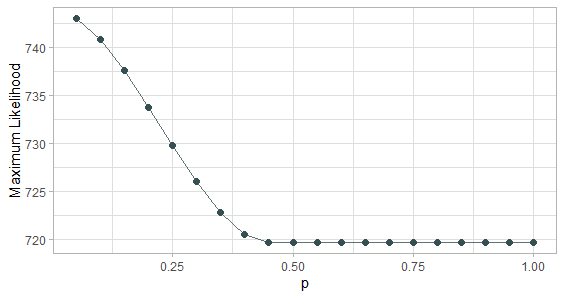

### figureA3.png

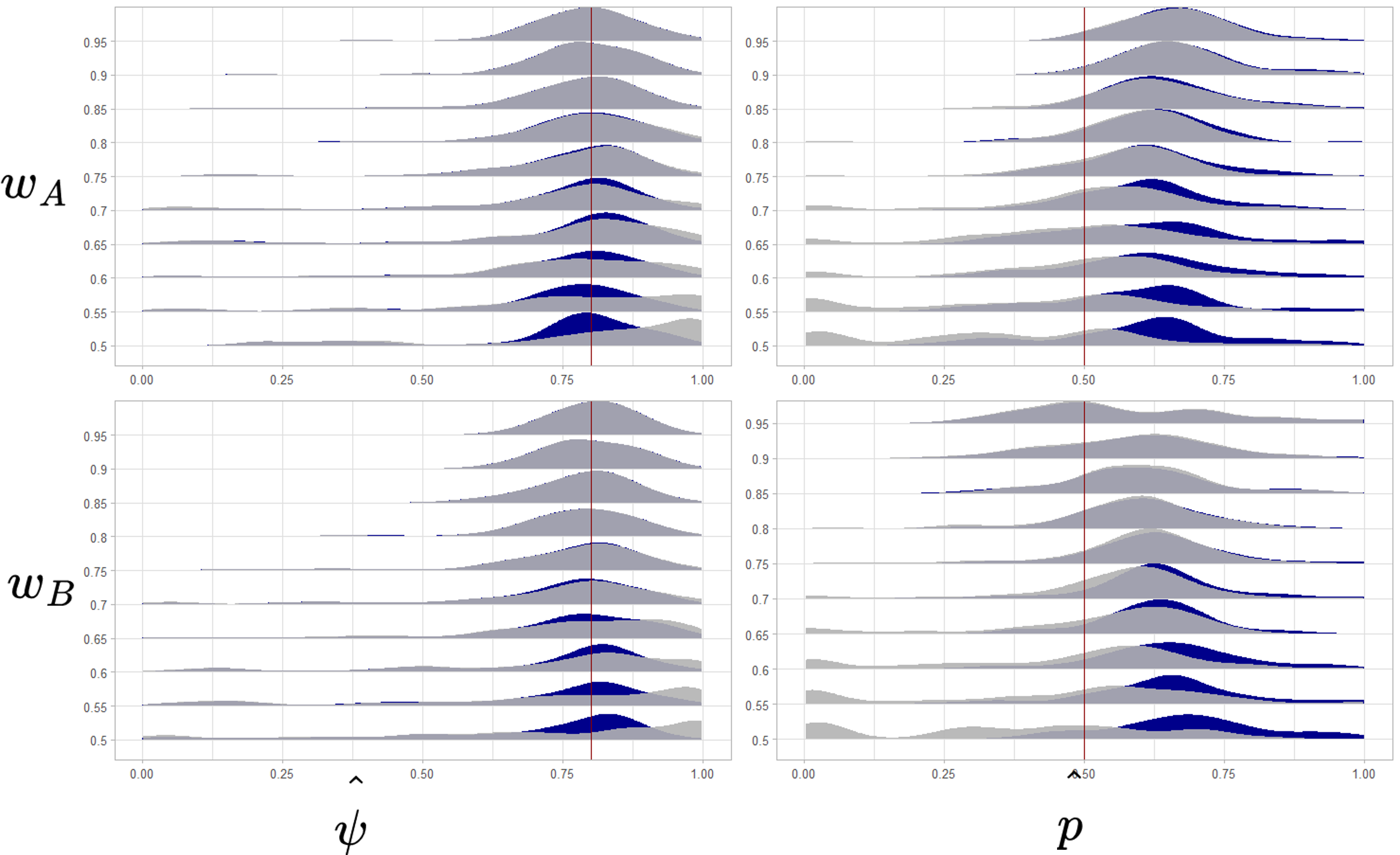

### figureA4.png

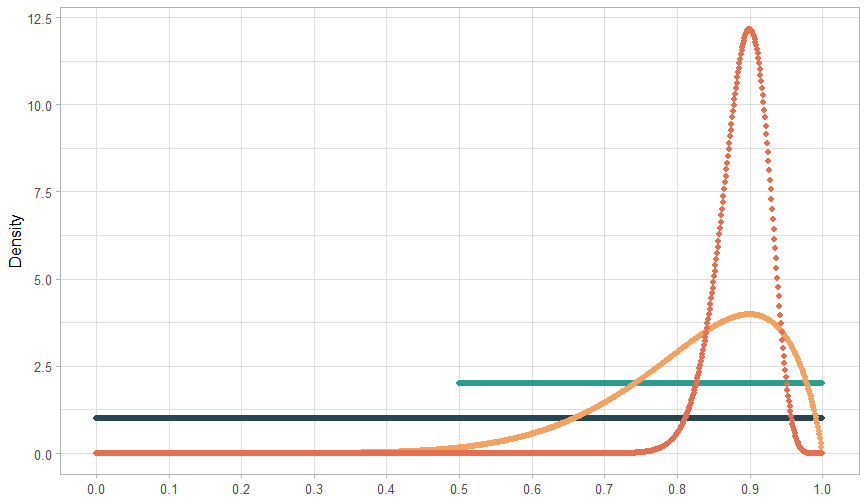

### figureA5.png

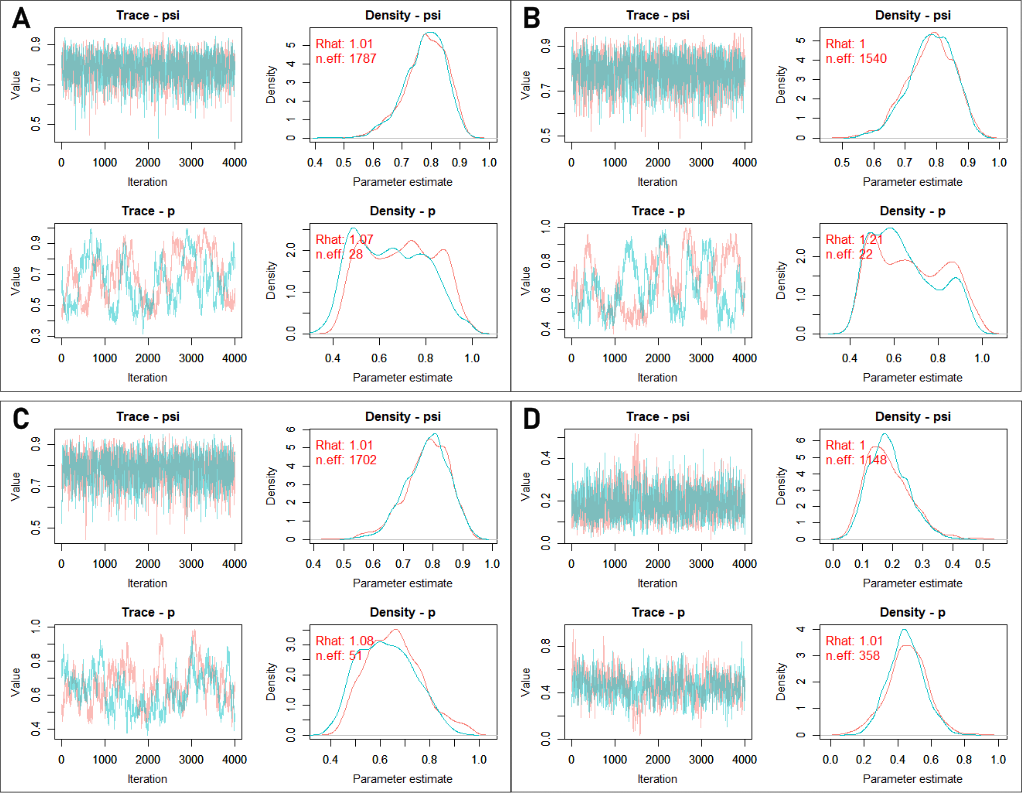

### figureA6.png

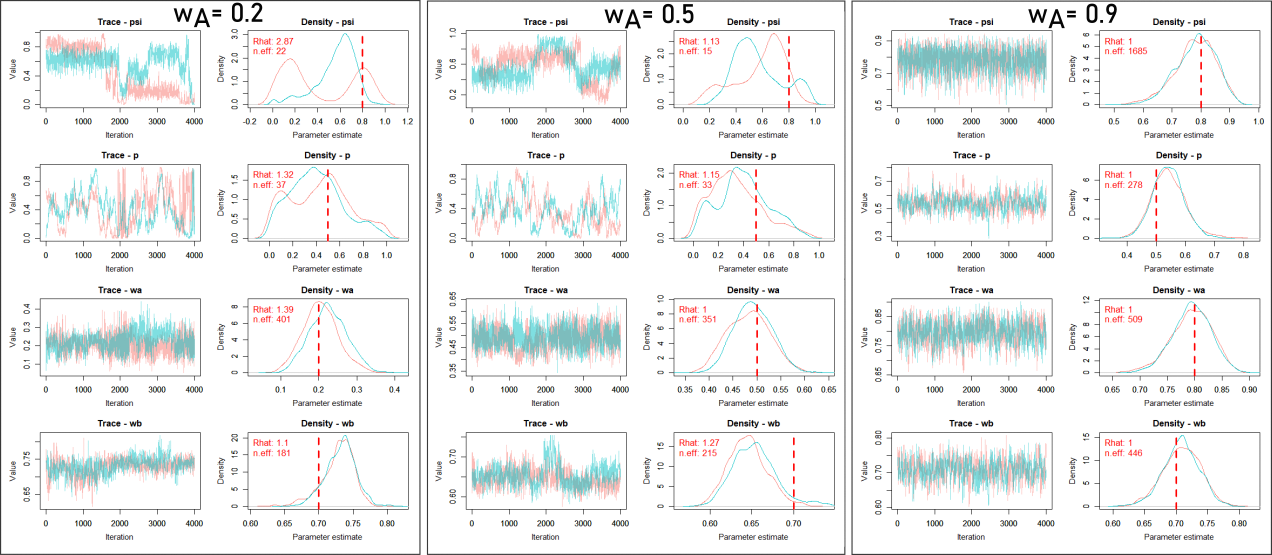
